# A plant single nucleotide polymorphism impacts nectar sugar composition, microbial diversity and pollinator visits

**DOI:** 10.64898/2026.04.09.717461

**Authors:** Guillaume Tueux, Nicolas Pouilly, Jordan Bernigaud-Samatan, Nicolas Blanchet, Marie-Claude Boniface, Olivier Catrice, Sébastien Carrère, Jérôme Gouzy, Marie-Pierre Jacquemot, Emmanuelle Lauber, Alexandra Legendre, Sandra Moreau, Marco Moroldo, Amandine Roldan, Aurélien Carlier, Nicolas B. Langlade

## Abstract

Nectar is a hub for plant-pollinator interactions, yet gene-level causal links between plant genetic variation, pollinator foraging and nectar microbial assembly remain poorly resolved. Using near-isogenic lines, innovative field time-lapse monitoring of pollinator visits and long-read amplicon sequencing of nectar microbiota, we show that a natural single-nucleotide variant at a cell-wall invertase gene (*HaCWINV2*) controls sunflower nectar chemistry and influences both pollinators and microbes. Plants homozygous for a loss-of-function *HaCWINV2* allele produce sucrose-rich nectar, resulting in fewer bee visits under field conditions, while bumblebee visitation remained unaffected. In pollinator-excluded flowers, invertase-deficient plants harboured greater fungal diversity and compositionally distinct communities, indicating that nectar sugar profiles act as ecological filters shaping the nectar microbiome. This loss-of-function allele was found rarely and only at the heterozygote state in wild sunflowers and was fixed in 35% of cultivated lines indicating a positive selection during domestication. Our findings establish a causal link between a single gene and nectar chemistry with cascading ecological effects in a plant–pollinator system, illustrating how subtle genetic changes scale up to alter nectar traits, microbial assembly and pollinator foraging behaviour.

## Introduction

Nectar is the main floral reward exchanged for animal pollination, and its chemistry is a central cue in plant–pollinator communication. Nectar sugars include sucrose, glucose and fructose, and their ratios may vary substantially between individual plants. Sugar content and composition may further alter viscosity and osmotic conditions, with consequences for the energetic and sensory rewards received by foragers, in particular honey bees^1,2^.

In sunflower (*Helianthus annuus*), nectar traits co-vary with pollinator visitation and are directly relevant to crop pollination and seed production^3–5^. Crop floral traits that alter pollinator visitation and reward quality may therefore influence both seed production and pollinator nutrition^1,6^. Yet the genetic basis of nectar sugar composition and its consequences for pollinator behaviour remain poorly resolved. Responses to floral traits are not necessarily uniform across pollinator taxa, and honey bees and bumblebees may respond differently to the same nectar and floral traits^7,8^.

Nectar compositiofn and quantity result from coordinated carbon allocation, secretion and extracellular metabolism in nectaries^2^. Sucrose export to nectaries relies on SWEET transporters, like for instance *SWEET9*^9^. Sucrose may be hydrolysed to the hexoses glucose and fructose by nectary-expressed cell-wall invertases^10,11^. Variations in the quantities of sucrose and hexoses in nectar have been associated with different pollination syndromes, suggesting that nectar sugar composition may contribute to shaping pollinator preferences^12^. Nectar is also a microbial habitat, populated by fungi and bacteria. Growth of yeasts and bacteria shape nectar chemistry through metabolic activity and emission of volatile signals^13^. Microbial metabolism may in turn affect pollinator health by altering the nutritional quality and chemical landscape of nectar^14^. Although dispersal by flower visitors is a major contributor to the structure of nectar microbial communities, ecological filtering may also influence community composition and structure^15–18^. This raises the possibility that plant genotype-dependent assembly of nectar microbial communities may mediate genetic effects on pollinator attractiveness.

In Arabidopsis, *AtCWINV4* is required for nectar production, and contributes to the typically hexose-rich nectar of the Brassicaceae^10,11^. *AtCWINV4* belongs to the GH32 glycosyl hydrolase family, which includes invertases and related fructan-active enzymes, and is characterized by a conserved catalytic core and C-terminal β-sandwich domain^19^. GH32 enzymes feature three conserved motifs—WMNDPNG, EC, and RDP—each containing an acidic residue that is central to catalysis and substrate binding^20^. Specifically, the aspartic acid (D) residue of the WMNDPNG motif is the nucleophile and the glutamic acid (E) of the EC-motif is the acid/base catalyst^21^. Although the D residue of the RDP motif is not thought to be involved in the catalytic mechanism of sucrose hydrolysis, it probably serves a key role in substrate binding and stabilization^22,23^. In sunflower, a homolog of the Arabidopsis nectar invertase *AtCWINV4* has been identified and named *HaCWINV2*^24^. Despite these mechanistic insights, direct evidence on how natural variation in *HaCWINV2* affects nectar traits is lacking. This is because nectar composition and quantity often correlate with other floral traits in crops such as sunflower, and quantitative trait loci (QTL) influencing pollinator attractiveness encompass broad genomic regions with many linked candidate genes^5,25^. For example, several genetic studies have identified *HaCWINV2* as a possible contributor to nectar quantity and composition in sunflower, but its functional role has remained unresolved^24–26^.

Here, we identified a commonly occurring loss-of-function variant in the nectar-expressed *HaCWINV2* gene that is linked to the production of sucrose-rich nectar in some sunflower genotypes. Using field monitoring of pollinator visits and nectar microbiome profiling of near-isogenic lines, we uncovered a direct link between genetic control of nectar sucrose–hexose balance, pollinator attraction and nectar microbial community structure.

## Materials and Methods

### Sunflower culture and phenotyping

Plants were grown at the INRAE experimental station of Auzeville-Tolosane (France) during the 2023, 2024 and 2025 field seasons. Each environment was defined by the combination of year and location and was included in statistical analyses to account for environmental variation. Hereafter, each environment will be referred to as “trial”. Plants were deployed in three non-overlapping subsets: a set of unnetted plants used exclusively for pollinator-attractiveness monitoring, and two independent sets of netted plants used for floral trait measurements and for nectar microbiota, respectively. To exclude insect foragers from plants used for floral traits and microbiota, heads in these two subsets were enclosed in pollination bags (Vilutis and Co.) prior to flowering. Wingscapes TimelapseCam Pro cameras were installed facing the head of each plant prior to flowering initiation until the end of flowering. Images were captured at five-minute intervals throughout the entire flowering period.

### Floral Traits

Sampling was conducted on netted heads on multiple collection days during the flowering period. On each sampling day, nectar was harvested from 6–8 florets per head using calibrated 1-µL glass microcapillaries (Hirschmann, Germany), approximately 24 h after anthesis at pistillate stage (10:00–14:00 local time). Nectar volume was measured from each capillary, and soluble-sugar concentration (°Bx) was determined with a refractometer (Bellingham & Stanley, UK). Plant-level means were used for analysis. Sugar mass per floret was calculated with the formula detailed by Cruden and Hermann^27^:

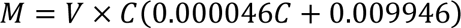

where M is the sugar mass in µg, V the nectar volume in nL and C the sugar concentration in g of sugar per 100 g of solution (% Brix). Concentrations of sucrose, glucose and fructose were determined enzymatically using the Megazyme K-SUFRG sucrose assay kit following the manufacturer’s instructions. Florets were collected after anthesis and preserved in 70% ethanol. Ten florets per sample were later randomly selected, scanned at 600 dpi using an Epson 11000XL scanner and measured for length and width using ImageJ^28^.

### Transcriptome analysis by RNA sequencing

We collected florets of plants of the inbred lines XRQ, IR and CI grown at the Heliaphen^29^ facilities in 2022. Florets at pistillate stage were collected between the daytime hours 10:00 and 12:00 and dissected under a stereomicroscope. Approximately 50 nectaries were collected per plant, frozen in liquid nitrogen and stored at -80°C before mRNA extraction with the RNAqueous Micro-kit (ThermoFisher AM1931). After verification of integrity by Agilent Bio-Analyzer 2100, RNA was sequenced by the genomics facility GetPlage (INRAE Auzeville-Tolosane). Data can be retrieved from the ENA SRA archive under study accession SRP451292. Paired-end sequences were pre-processed with the ‘nf-core/rnaseq’ v 3.0 pipeline (https://zenodo.org/records/4323183). Sequence pseudoalignment was then performed on the HanXRQr2.1 genome using salmon v 1.4.0^30^. Sequence counts were annotated at the gene level. Then, the counts were converted to ‘counts per million’ (CPM) and filtered by retaining only genes with CPM >0.5 in at least three libraries.

### Sunflower genome sequencing and analysis

The genome sequences of the XRQ and IR inbred lines, with accessions HanXRQr2.1-SUNRISE and HanIRr1.0-20201123 were downloaded from www.heliagene.org^31,32^. The genome of the CI inbred line was sequenced using a combination of PacBio CCS, with Bionano optical map and annotation using genotype-specific PacBio IsoSeq mRNA as previously described^31^ and with the support of the International Consortium for Sunflower Genomics. The annotated genome of the CI inbred line is available under the accession number HannCIr1.0-20220819 at www.heliagene.org.

The analysis of genome sequencing datasets of wild and cultivated sunflower is described in Supplementary Methods. Variant tables limited to the *HaCWINV2* locus were extracted from whole genome variant tables format with bcftools^33^, with -r = HanXRQChr01:64034704-64043774.

### Identification of CWINV homologs in *H. annuus* genome sequences

Homologs to cell wall invertase proteins were identified with tBLASTn, using the amino acid sequence of AtCWINV4 of *A. thaliana* (TAIR accession number AT3G52600). Protein sequences from HaCWINV family members were aligned using ClustalW as implemented in the R package *msa* (v1.42.0), with default parameters for amino acid sequences. Pairwise distances based on sequence identity were computed with the R package *seqinr* (v4.2-36). A phylogenetic tree was then reconstructed from this distance matrix using the neighbor-joining algorithm implemented in the R package *ape* (v5.8-1)^34^.

Amino acid sequences of experimentally characterized GH32 family proteins, including AtCWINV4, were downloaded from the RCSB PDB database. Predicted protein sequences of HaCWINV2 homologs were extracted from the genome sequences of *Helianthus annuus* cultivars XRQ, IR and CI. Amino acid sequences were aligned using MAFFT v7^35^ in auto mode.

### Strains and media

*Komagataella phaffii* strain OPENPichia *hoc1^tr^*and derivatives were cultured at 28°C in yeast extract peptone dextrose (YPD; 1% yeast extract, 2% peptone and 2% D-glucose), BMY (1% yeast extract, 2% peptone, 1.34% yeast nitrogen base without amino acids, 100 mM potassium phosphate buffer pH 6 and 4.10-5% Biotin), buffered methanol-complex medium (BMMY; BMY with 0.5% v/v methanol), or buffered glycerol-complex medium (BMGY; BMY with 1% glycerol) as previously described^36^. Media were supplemented with hygromycin B (100 µg/mL) as appropriate. *Escherichia coli* TG1 was grown in Luria-Bertani (LB) medium at 37°C, with or without hygromycin B 100 µg/mL.

### *K. phaffii* transformation

A synthetic gene fragment of *HaCWINV2* was designed based on the amino acid sequence of the HanXRQr2_Chr01g0013971 gene (Genbank accession KAF5821434). The first 31 AAs of the protein were not included in the synthetic gene construct to remove possible interference from a putative plant secretion signal peptide. A consensus MFα PrePro signal peptide was fused to the N-terminus of the protein sequence, and an 8xHis tag at the C-terminus. The resulting fragment was synthesized by Twist Bioscience (San Francisco, USA) after codon optimization for *K. phaffii* using the company’s online tool. The resulting synthetic gene fragment was assembled with OpenPICHIA parts harbouring a *PAOX1* promoter derived from the pPTK001_P2_pAOX1 plasmid, a fragment harbouring an *AOX1* transcription terminator, and cloned into the pPTK196_P8_HygR plasmid by Golden Gate assembly with BsaI restriction enzymes, yielding plasmid pEL226. A variant was synthesized following an identical strategy, in which the AGA codon coding for Arg 191 of HanXRQr2_Chr01g0013971 was replaced by a TGT codon to yield construct *HaCWINV2_XRQ_R191C_* harboured on plasmid pEL225. Plasmids were fully sequenced in-house using ONT Nanopore technology as described elsewhere (https://hal.science/hal-04684858v1). Nucleotide sequences of pEL225 and pEL226 were deposited in NCBI Genbank, with accession numbers PX504709 and PX504708, respectively. Plasmid DNA extracts of pEL225 and pEL226 were linearized by digestion with SacI. Synthetic gene constructs were introduced into *K. phaffii* OPENPichia *hoc1^tr^*by electroporation following established protocols^37^. Transformants were isolated on YPD medium supplemented with hygromycin B and stored at -80 °C in 25% glycerol after passaging once on YPD. The presence of the insert was assessed by colony PCR using primers CGACGCTCATCTGCATGTG and AGGCTGATCAGGAGCAAGC. Six clones per construct were selected and their genomes fully sequenced.

### *K. phaffii* genome sequencing and mapping of insertion site

*K. phaffii* genomic DNA was isolated from overnight cultures in BMGY medium and genomic DNA was isolated using the MasterPure DNA purification kit (Epicentre Reference). DNA library preparation was carried out with the Oxford Nanopore rapid barcoding 96 kit v114 and sequenced using an Oxford Nanopore P2 solo instrument equipped with an R10.4.1 flow cell. Basecalling was performed with dorado v0.9^38^ in GPU mode on a Linux PC equipped with a, NVIDIA A6000 graphics card and 48 Gb of on-board memory. Reads were filtered with chopper^39^ with the following settings: -q 10 -l 1500 --headcrop 10 --tailcrop 10. Filtered reads were mapped against the reference genome of *K. phaffii* NRRL Y-11430 (Genbank accession GCA_030557435.1) using the Clair3 program with following settings: --platform=ont, model = r1041_e82_400bps_sup_v430, --min_coverage=5, --enable_long_indels, --include_all_ctgs, and --haploid_precise^40^. Sequencing data were deposited at the European Nucleotide Archive, with accession numbers ERS27228307-ERS27228318.

### Recombinant protein expression and enzymatic activity assay

Overnight cultures of *K. phaffii* strains were prepared in BMGY medium. Cultures were centrifuged at 3,000 x g for 5 min and washed once in fresh BMMY medium. Pellets were resuspended in 5 mL of BMMY medium supplemented with 1 g/L of sucrose to an OD600nm = 0.8 and cultured for 72 hours at 28°C with shaking. An additional 0.5% (v/v) methanol was added to the culture every 24 hours. Culture supernatants were collected by centrifugation at 10,000 x g for 4 min and analysed by SDS-PAGE. Although the yields were generally low and prevented the purification of His-tagged proteins from supernatants, six clones for each construct were selected for further experiments based on similar expression profiles of the recombinant protein of MW = ca. 72.9 kDa. Two out of 6 clones for HaCWINV2-XRQ_R191C_ were excluded because they carried multiple tandem transgene insertions (ENA accessions ERS27228316 and ERS27228317). The final concentration of sucrose in the supernatant of the selected clones was determined using the Megazyme K-SUFRG sucrose assay kit following the manufacturer’s instructions.

### Development of Near Isogenic Line pairs for *HaCWINV2* alleles

Two inbred lines, with CI as the female parent and IR as the male parent were crossed to produce F1 hybrids, then self-pollinated at F2 and later stages. A subset of 95 F3 seedlings were screened using a PCR Allelic Competitive Extension genotyping essay for a SNP located within the *HaCWIN2* gene (HanXRQr2_Chr01g0013971) (Supplementary Table 1). For this SNP, two allele-specific primers and one flanking primer were designed using BatchPrimer3^41^. The genotyping was performed using the PACE 2.0 Genotyping Master Mix (3CR Bioscience, Essex, UK). Thermal cycling and fluorescent signal detection were conducted on a Roche LightCycler 480-II System, following the manufacturer’s instructions. Four pairs of Near Isogenic Lines per F2 family were thus identified.

### Axiom genotyping of NIL pairs

Genomic DNA of 12 plants from each line was extracted using previously described protocols^42^. Genotyping using pooled DNA of the NILs was performed at the Gentyane platform (UMR INRAE, Clermont-Ferrand, France) using the Axiom Multispecies 9-Genotyping Array (384HT-array, Cat. No. 551416, Life Technologies), which includes 10,763 sunflower SNP markers. Genotypic data were obtained with Axiom Analysis Suite v5.2 (Life Technologies) and visualized with Flapjack^43^.

Based on Axiom genotyping, the CI and IR parent inbred lines shared 69.3% of markers. Among the four NIL pairs, Pair 1 and Pair 2 showed the highest genome-wide similarity (86.1% and 92.0% of shared SNP markers between lines of a pair differing at the *HaCWINV2* locus, respectively). Lines of Pair 3 and Pair 4 were more divergent with 82.0% and 75.9% of shared markers between lines of a pair, respectively. Because of a technical issue with propagation for Pair 3 and higher genetic divergence, we restricted our phenotyping experiments to plants belonging to Pair 1 and Pair 2.

### Analysis of floral traits

Plant-level means were calculated by averaging nectar trait measurements across the sampled florets for each plant. We required measurements from at least three florets per plant; plants not meeting this criterion were excluded. Each NIL pair was analysed separately (Pair 1: n = 33 plants; Pair 2: n = 40 plants). Within each pair, nectar volume (µL per floret) and sugar mass (µg per floret) were analysed by including the *HaCWINV2* allele and the trial as fixed effects in the models. Model assumptions were evaluated by inspecting residual Q–Q plots, testing residual normality (Shapiro–Wilk^44^), and assessing homogeneity of variances across alleles (Levene’s test). Full diagnostic results are reported in Supplementary Table 2. According to the Shapiro–Wilk^44^ test, sugar mass in Pair 2 showed a slight deviation from normality. However, inspection of the corresponding residual Q–Q plot (Supplementary Fig. 1) did not reveal any substantial departure from normality. Allele effects were assessed using within-pair estimated marginal mean contrasts (emmeans^45^), and averaged over trials. All tests were two-sided. The capillary-level measurements supporting these plant-level summaries are provided in Supplementary Data.

Floret length and width (mm) were analysed at the plant level (n = 24 plants; 12 plants per NIL pair) and compared between *HaCWINV2* alleles within each pair using two-sided Welch t-tests^46^. Full assumption checks and test outputs are provided in Supplementary Table 3. The raw floret-level measurements used for these analyses are provided in Supplementary Data.

### Quantification of pollinator visits

Insect visitation was quantified from time-lapse images acquired every 5 min throughout the experiment. We used a detection model based on the YOLO11x algorithm (https://github.com/ultralytics/ultralytics), developed internally and available at https://forge.inrae.fr/astr/public/pollicrop to detect and classify insects into two categories:

(i) bees (including *Apis mellifera*., *Megachile* spp., *Halictidae*) and (ii) bumblebees. Model performance was validated by human expert annotation and evaluation of a subset of 5,000 randomly selected images. The detection model demonstrated the following performance metrics on the validation dataset: precision 0.906, recall 0.954, and F1-score 0.929 for bees; and precision 0.750, recall 1.000, and F1-score 0.857 for bumblebees. Further details on model training, dataset construction, and augmentation strategies are provided in Supplementary Methods.

Meteorological data were collected hourly by a weather station (INRAE 31035002) located within 500 m from the field, and data were extracted via the Climatik interface (INRAE; https://agroclim.inrae.fr/climatik/). Each image was time-stamped and linked to the corresponding hourly meteorological record. We aggregated detections at the plant × hour scale by summing counts across all images taken from the same plant within the same clock-hour on the same date (Europe/Paris time). This aggregation, together with plant-level random intercepts and meteorological covariates, mitigates temporal auto-correlation. The aggregated plant × hour dataset and associated meteorological covariates are provided in Supplementary Data. Analyses were restricted to daytime (06:00–21:59), resulting in a final dataset of n = 63 plants. For each plant separately, observations were retained up to the end of flowering. This date was determined using bee visits, i.e. by the time at which cumulative pollinator visits reached 99% of the total over the camera acquisition period. This trimming was necessary to remove late-flowering periods with very few remaining florets (< 10).

Bee and bumblebee counts were analysed separately using negative-binomial generalized linear mixed models (GLMMs; NB2) fitted with glmmTMB^47^ (log link) with *HaCWINV2* allele, NIL pair and their interaction included as fixed effects. Negative binomial models were chosen because Poisson models showed poorer fit, with substantially higher Akaike Information Criterion (AIC) values. Let *y_it_* denote the aggregated count for plant *i* in time unit *t* (date × clock-hour), and *N*_*it*_ the number of images contributing to that unit. Because meteorological covariates were available at the same hourly resolution as the response and detections were aggregated at the plant × hour level, we did not include additional date- or hour-of-day effects in the primary models to avoid redundancy with weather predictors, and repeated measures were handled via plant-level random intercepts. We assumed:

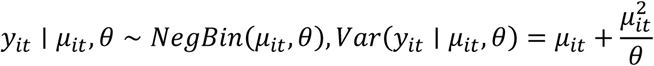

μ*_it_* = 𝔼[*y_it_*] denotes the mean (expected) visitation count for plant *i* at time *t*, and θ is the negative-binomial dispersion (size) parameter controlling overdispersion. The mean was modelled with a log link and an offset for sampling effort:

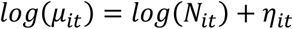

so that *exp*(η*_it_*) represents the expected visitation rate per photograph (i.e. *µ_it_*/**N*_it_*). The linear predictor was:

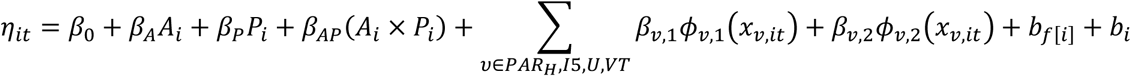

where *A_i_* encodes the *HaCWINV2* allele with *HaCWINV2+* as the reference level (*A*_i_ = 0 for *HaCWINV2+* and *A_i_* = 1 for *HaCWINV2-*). While *P*_*i*_ encodes the NIL pair with Pair 1 as the reference level (*P_i_* = 0 for Pair 1 and *P_i_* = 1 for Pair 2). The terms *x_v,it_* are hourly meteorological covariates (*PAR_H_* (hourly photosynthetically active radiation, *J*. *cm*^-2^), *I*5 (actinothermic index at 50 cm, °C), *U* (hourly relative humidity, %) and *VT* (hourly wind-run distance, km). Meteorological covariates were selected based on their documented influence on pollinator flight and foraging activity in bees and other pollinators^48^. To accommodate potentially non-linear relationships between visitation rates and these environmental drivers, each covariate was modelled using a second-order orthogonal polynomial basis (ϕ*_v_*_,1_, ϕ*_v_*_,2_), as implemented in R with *poly*(*x*, 2, *raw* = *FAKSE*) from the stats package. Random intercepts accounted for repeated observations within trials and plant identity, with 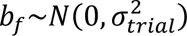 and 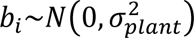.

Under this coding, *exp*(β*_A_*) represents the effect of the *HaCWINV2−* allele relative to *HaCWINV2+* in the reference NIL pair, whereas β_AP_ represents the difference in the allele effect between NIL pairs. To facilitate interpretation, marginal means and contrasts were estimated with emmeans^45^ on the response scale, both as an overall allele effect averaged across NIL pairs and as pair-specific allele contrasts.

Model adequacy was assessed using DHARMa^49^ (10,000 simulations). Dispersion and zero-inflation tests provided no evidence for over or under-dispersion or excess zeros across models. KS tests of residual uniformity were significant in one model, which can occur with large sample sizes and may reflect minor departures from the assumed mean–variance structure. Inspection of DHARMa residual plots did not indicate strong patterns (Supplementary Fig. 2). Bootstrapped outlier tests did not indicate a significant excess of outliers in either model. Multicollinearity and singularity were checked with the performance R package and indicated low collinearity (maximum VIF < 5) and no singular fits. Full diagnostics are reported in Supplementary Table 4.

### PCR and sequencing of nectar microbiota samples

Nectar microbiota was characterized on netted plants located in the same fields as those used for floral trait measurements. Nectar was collected in 2023 and 2024, 24 h post-anthesis at the pistillate stage, using sterile 1 µL or 5 µL glass microcapillaries (Hirschmann). Nitrile gloves disinfected with 70 % ethanol were used and cleaned between plants to prevent cross-contamination. We included controls consisting of empty capillaries that we waved in the air close to the inflorescences and touching its tip with gloved fingers. These field controls were taken at the start and end of each sampling session (four in 2023, five in 2024). Capillaries were immediately stored in sterile 1.5 mL centrifuge tubes and kept at −80 °C until analysis. DNA was extracted with the DNeasy PowerSoil Pro kit (QIAGEN). All samples, including controls, were processed in parallel, and a positive control (ZymoBIOMICS Microbial Community Standard, Zymo Research) was included and carried through DNA extraction, library preparation, sequencing, and the bioinformatic pipeline. Nectar microbial culturability was assessed in 2024 by plating nectar samples on semi-selective media. Nectar was diluted in 500 µL sterile phosphate-buffered saline (PBS), spread-plated onto tryptic soy agar (TSA) and R2A agar plates using sterile beads, and incubated for minimum 3 days at 28°C. Colony-forming units (CFUs) were then counted for each plate. The near full-length bacterial 16S-rRNA gene and fungal ITS1-ITS4 region were amplified with adapter-tailed primers 16SF/16SR and ITS1F/ITS4R, respectively (Supplementary Table 5). PCR was performed using Platinum™ Taq DNA Polymerase No-DNA (Thermo Fisher Scientific). Amplicons were barcoded with the Nanopore EXP-PBC096 and libraries prepared with SQK-LSK114 (Oxford Nanopore Technologies) using the protocols from the manufacturer. Sequencing was performed on a PromethION 2 platform (Oxford Nanopore Technologies) using R10.4.1 flow cells. Raw nanopore signals in POD5 format were basecalled with Dorado v0.9 (Oxford Nanopore Technologies) using the dna_r10.4.1_e8.2_400bps_sup@v5.0.0 model and the --no-trim and --barcode-both-ends parameters. Sequencing reads per sample were adapter-trimmed, oriented, and filtered using Cutadapt^50^, chopper^39^ and custom Python scripts deposited at https://forge.inrae.fr/aurelien.carlier/attracthol. Briefly, sequences were first oriented by searching for the 1st round PCR primer sequences on the forward and reverse-complemented sequence for each read. PCR primer and sequencing adapter sequences were removed with cutadapt. Reads with an average Phred score < 20 were discarded, as well as reads whose length fell outside of the ranges of 1300 - 1600 bp for 16S rRNA amplicons and 300 - 1000 bp for ITS amplicons. When applicable, read sets were randomly subsampled to 50 000 sequences using the sample function of seqtk (https://github.com/lh3/seqtk).

Sequences were clustered into OTUs at 97% identity with VSEARCH^51^. OTUs corresponding to chimeric and sequencing artefacts were removed with the VSEARCH UChime implementation and the Mumu software (https://github.com/frederic-mahe/mumu) using default settings. The resulting OTU tables were imported in the Qiime 2 software v.2023.9.1^52^ or taxonomic assignment using the VSEARCH classifier with a 80% sequence identity threshold and the Silva 138 NR99 database^53^ for 16S rRNA sequences, or the UNITE NR 99^54^ database for fungal ITS sequences. All raw 16S rRNA and ITS sequencing data, including nectar samples, negative controls and mock community dilutions, have been deposited in the European Nucleotide Archive under study accession PRJEB106052.

### Microbial diversity analysis

All computations were carried out in R using phyloseq^55^, vegan^56^, ape^34^, iNEXT^57^ and emmeans^45^. OTU tables were filtered using blanks and empty-capillary controls: OTUs reaching > 5% relative abundance in any negative-control library (within-control normalization) were removed from the entire dataset. OTUs of non-microbial origin (non-fungal ITS sequences or chloroplast) were discarded. In addition, OTUs assigned to the class Malasseziomycetes, considered to be of human origin, were removed. We then applied iterative abundance and depth filtering until convergence by removing OTUs with < 10 total reads and samples with < 1,000 reads. Rarefaction curves were generated on the filtered ITS dataset (post-contaminant and post-depth filtering) to verify that the remaining sequencing depth was sufficient to capture fungal diversity (Supplementary Fig. 3). Alpha diversity was calculated as coverage-standardized Hill numbers^58^ at 95% sample coverage using iNEXT^57^. We report richness (Hill q = 0), the exponential of Shannon entropy (Hill q = 1; effective number of common OTUs, hereafter ENS1), and the inverse Simpson (Hill q = 2; effective number of dominant OTUs, hereafter ENS2). Richness was square-root transformed prior to analysis, whereas ENS1 and ENS2 were analysed on the original scale. Model assumptions were assessed on the fitted ANOVA models: normality of residuals was tested with Shapiro–Wilk tests^44^, and homoscedasticity was evaluated using Levene’s tests across *HaCWINV2* allele × NIL pair groups. Full assumption checks and test outputs are provided in Supplementary Table 6. Each metric was analysed using an ANOVA model with fixed effects for the *HaCWINV2* allele, the NIL pair, their interaction and the trial year. Post-hoc comparisons used estimated marginal means focused on the *HaCWINV2* allele main effect. Community structure was quantified using Bray–Curtis distances computed from relative abundances, Jaccard distances computed on presence/absence data, and unweighted UniFrac distances computed from raw counts using a rooted phylogeny. Homogeneity of multivariate dispersions was assessed for *HaCWINV2* allele, NIL pair, and trial year with betadisper followed by permutest (9,999 permutations) for each distance. Effects of *HaCWINV2* allele, NIL pair and trial year were then tested by PERMANOVA (99,999 permutations, marginal tests, by = margin). Dispersion tests indicated no evidence for heterogeneity of dispersions with respect to *HaCWINV2* allele, NIL pair or trial year for Bray–Curtis or Jaccard distances. Unweighted UniFrac results were not interpreted further because multivariate dispersions differed significantly between lines (Supplementary Table 7). OTUs were classified as shared if they were detected (raw count > 0) in at least one *HaCWINV2−* sample and at least one *HaCWINV2+* sample. OTUs detected exclusively in samples from a single allele group were classified as *HaCWINV2−* only or *HaCWINV2+* only. For each sample, we quantified the fraction of reads assigned to allele-specific OTUs by summing counts across OTUs in the corresponding “only” category and dividing by the total read count of that sample; proportions were then averaged within each allele group. The R codes used for this analysis are found at https://forge.inrae.fr/aurelien.carlier/attracthol.

### Estimation of the microbial detection limit by amplicon sequencing

To estimate the 16S rRNA–amplicon limit of detection in nectar, we generated a 10-point serial dilution series of the community standard in a pooled nectar DNA sample composed of 10 plants collected in 2022 under the same protocol, used to develop the workflow and yielding community profiles similar to those from the 2023 and 2024 trials. The mock input was set to 2×10^9^ cells mL^−1^ (from the manufacturer’s and assuming full recovery). Dilution 1 mixed 2 µL mock DNA with 18 µL nectar DNA (1:10; 4.0×10^6^cells). Dilutions 2–10 were successive 6.5-fold serial dilutions. PCR amplification of 16S amplicons, sequencing by ONT Nanopore technology, and analysis were realized as described in the section PCR and sequencing of nectar microbiota samples. We defined “MOCK OTUs” as all OTUs present in the pure MOCK sample and, for each dilution and nectar-only sample, we calculated the proportion of reads assigned to MOCK OTUs and the percentage of reads assigned to non-chloroplast OTUs. Theoretical input cell numbers were derived from the manufacturer’s nominal concentration (2×10⁹ cells mL⁻¹), and the dilution factor of each sample. The nectar-only sample provided the background level of non-chloroplast reads against which the dilution series was compared to infer a practical detection limit.

### Statistical software and packages

All analyses were performed in R (v4.5.1) using RStudio (v2023.9.1.494). The following R packages were used: *ggplot2* (v4.0.1), *tidyr* (v1.3.1), *dplyr* (v1.1.4), *readxl* (v1.4.5), *phyloseq*^55^ (v1.52.0), *ape*^34^ (v5.8.1), *iNEXT*^57^ (v3.0.1), *tibble* (v3.3.0), *vegan* (v2.7.1), *car*^59^ (v3.1.3), *emmeans*^45^ (v2.0.0), *purrr* (v1.0.4), *scales* (v1.4.0), *forcats* (v1.0.1, *glmmTMB*^47^ (v1.1.13), *DHARMa*^49^ (v0.4.7), *performance*^60^ (v0.15.2).

## Results

### Polymorphism in the *HaCWINV2* coding sequence correlates with nectar sugar composition

During field trials, we observed that three sunflower inbred lines (IR, CI and XRQ) displayed marked differences in nectar sugar composition (Supplementary Fig. 4). The IR and XRQ lines produced nectar that contained almost exclusively hexoses (96.2% ± 1.2% and 95.3% ± 6.5% hexoses, mean ± SE, respectively), whereas nectar from the CI line showed substantially higher sucrose content (29.3% ± 15.0% sucrose, mean ± SE). Previous reports indicated that cell-wall invertase enzymes expressed in the nectary may be involved in controlling nectar sugar composition. The genome of *Helianthus annuus* XRQ contains six genes coding for putative *A. thaliana* CWINV homologs, only one of which (HanXRQ2 Chr01g0013971, later named *HaCWINV2*) is expressed in the nectary (Fig. 1A). Alignment of the predicted protein sequences of HaCWINV2-XRQ, HaCWINV2-IR and HaCWINV2-CI revealed polymorphisms resulting in five amino-acid changes, including two in a putative secretion signal (Supplementary Fig. 5). One polymorphism modified the RDP motif, conserved in proteins of the GH32 family, to a CDP sequence in the CI line at position 191 (Fig. 1B). Although not identified as directly involved in substrate binding or catalysis, the modification of this residue in the conserved RDP motif may affect the ability of enzyme to convert sucrose into hexoses.

**Figure 1.**
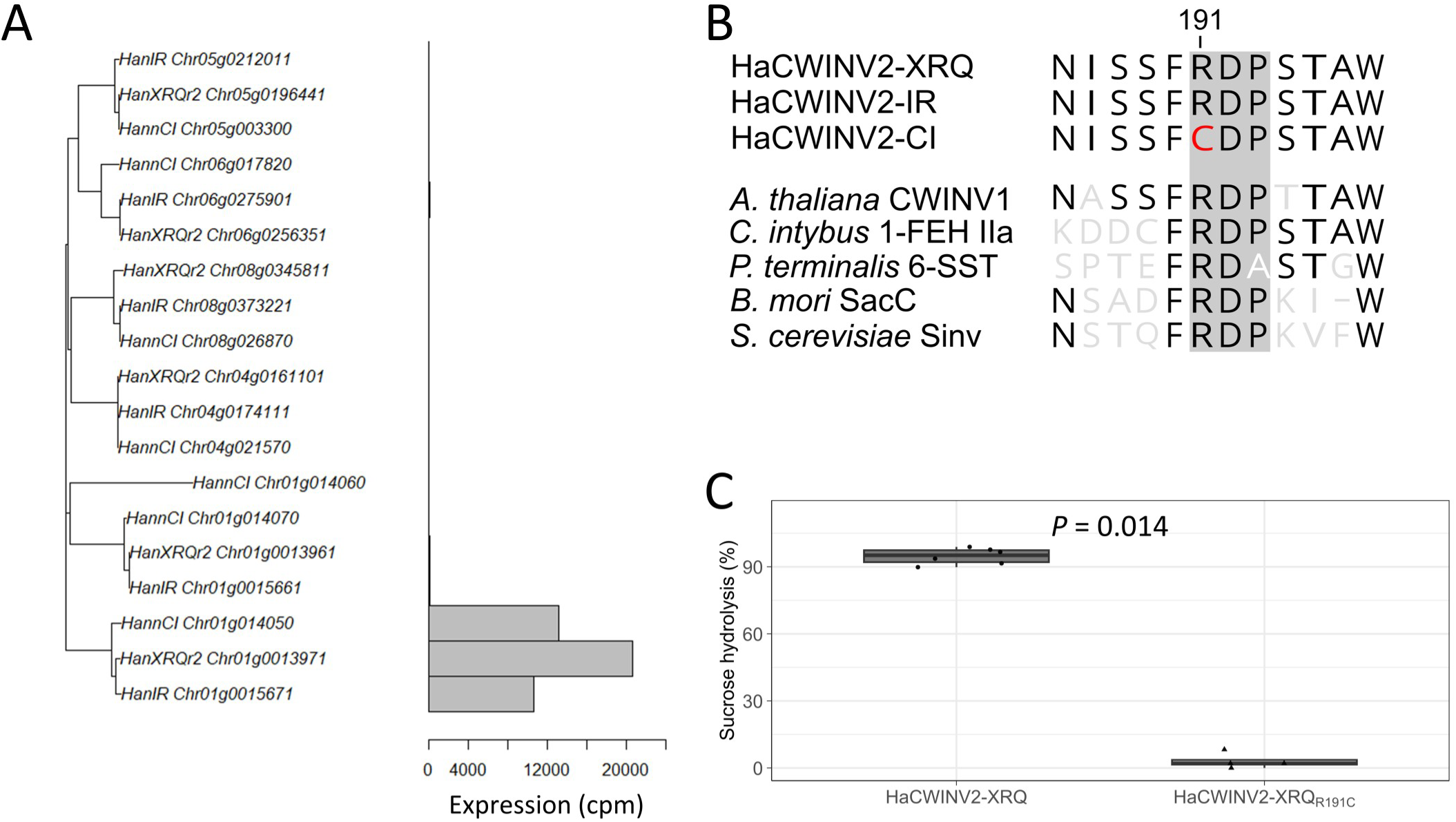
HaCWINV2 is a sucrose invertase expressed in the nectary. A. Phylogenetic tree of sunflower candidate cell-wall invertase. Prefixes HanXRQr2, HannCI and HanIR correspond to the locus tags of lines XRQ, CI and IR, respectively. Horizontal bars indicate gene expression (CPM) derived from RNA-seq data obtained from dissected nectary tissue. Locus tag *HanXRQr2_Chr01g0013971* correspond to the *HaCWINV2* gene in the XRQ reference genome. B. Partial alignment of HaCWINV2 predicted protein coding sequence of sunflower lines XRQ, CI and IR, as well as representative eukaryotic members of the GH32 family (full alignment in Supplementary Fig. 5). Conserved residues relative to the HaCWINV2-XRQ sequence are depicted in black, other residues in grey or white. The conserved RDP motif is highlighted in grey, and a cysteine replacing the conserved arginine of the RDP motif in the HaCWINV2-CI sequence is shown in red. C. Sucrose hydrolysis in *K. phaffi* cultures expressing HaCWINV2 variants. Sucrose (1 g/L) was added to cultures of at least four independent clones for each construct. Remaining sucrose in the supernatant was measured independently three times after 72h of incubation. Values correspond to the difference between initial sucrose concentration and remaining sucrose concentration, expressed as percentages. Each point represents one independent clone, calculated as the mean of three technical replicates. *P* values were calculated using a two-sided Wilcoxon rank-sum test on clone means.

### The HaCWINV2 R191C mutation results in a loss of sucrose hydrolysis activity

To investigate how a change of residue from Arginine(R) to Cysteine(C) at position 191 of HaCWINV2 affects enzymatic activity, we designed a synthetic gene construct modelled on the *HaCWINV2-XRQ* allele fused to a yeast secretion signal sequence, together with a variant differing only in an Arg or Cys residue at position 191. We expressed the synthetic constructs in the yeast *Komagataella phaffii* (*K. phaffii*), which lacks an endogenous sucrose hydrolase^61^. Cultures expressing the *HaCWINV2-XRQ* allele depleted exogenously-supplied sucrose. In contrast, cultures expressing the HaCWINV*2*-XRQ_R191*C*_ variant lacked detectable sucrose hydrolysis activity (Fig. 1C), despite secreting similar quantities of the protein (Supplementary Fig. 6). This indicates that loss of enzymatic function, rather than difference in expression, is responsible for the inability of the HaCWINV2-XRQ_R191C_ variant to hydrolyse sucrose. This shows that the genome of the CI line encodes a catalytically impaired version of the nectary-expressed cell wall invertase HaCWINV2.

### Frequency of loss of function mutations in cultivated and wild sunflower lines

We screened a panel of 419 cultivated sunflower lines to determine the frequency of the HanXRQChr01:64037184 G→A polymorphism, which results in the R→C residue substitution at position 191 (Supplementary Data). In this dataset, 270 lines were homozygous for the wild-type (G) allele (64.4%), 3 were heterozygous (0.7%) and 146 were homozygous for the loss-of-function (A) allele (34.8%). We also analysed a dataset of 184 wild sunflower accessions, of which 177 were homozygous for the wild-type allele (96.7%), 6 were heterozygous (3.3%), one did not have a call at that position, and none (0%) was homozygous for the loss-of-function allele (Supplementary Data). These results show that the allele corresponding to catalytically impaired forms of HaCWINV2 is frequent in cultivated lines, but rare in wild accessions.

### *HaCWINV2* controls nectar sugar composition

To confirm that polymorphism at the *HaCWINV2* locus is directly responsible for the variable sucrose/hexose ratio in sunflower genotypes, we generated near-isogenic line (NIL) pairs by repeated crossing of the CI and IR lines. Because targeted mutagenesis approaches are not yet routinely available in sunflower, this strategy allowed us to isolate the effect of the *HaCWINV2* locus while minimizing differences in genetic background. We selected two NIL pairs whose members differed primarily at the *HaCWINV2* locus. The two lines of Pair 1 shared on average 86% of markers, while the two lines of Pair 2 shared 92%. For clarity, we will refer from this point on to lines of each pair harbouring the functional IR allele as *HaCWINV2+* and the CI allele as *HaCWINV2-,* respectively.

For both NIL pairs, the proportion of sucrose in nectar varied between the two *HaCWINV2* alleles (Fig. 2A). Nectar of *HaCWINV2+* plants contained ∼1% sucrose (Pair 1: mean = 0.8%; Pair 2: mean = 1.1%), whereas the nectar of *HaCWINV2−* plants consistently showed much higher sucrose proportions (Pair 1: 24.3%; Pair 2: 34.5%). This difference was statistically significant for each NIL pair (Welch’s t-test: Pair 1, P = 0.026; Pair 2, P = 0.009). These data confirm that the CI sunflower line expresses a catalytically impaired variant of *HaCWINV2*, which is responsible for the elevated sucrose content in nectar.

**Figure 2:**
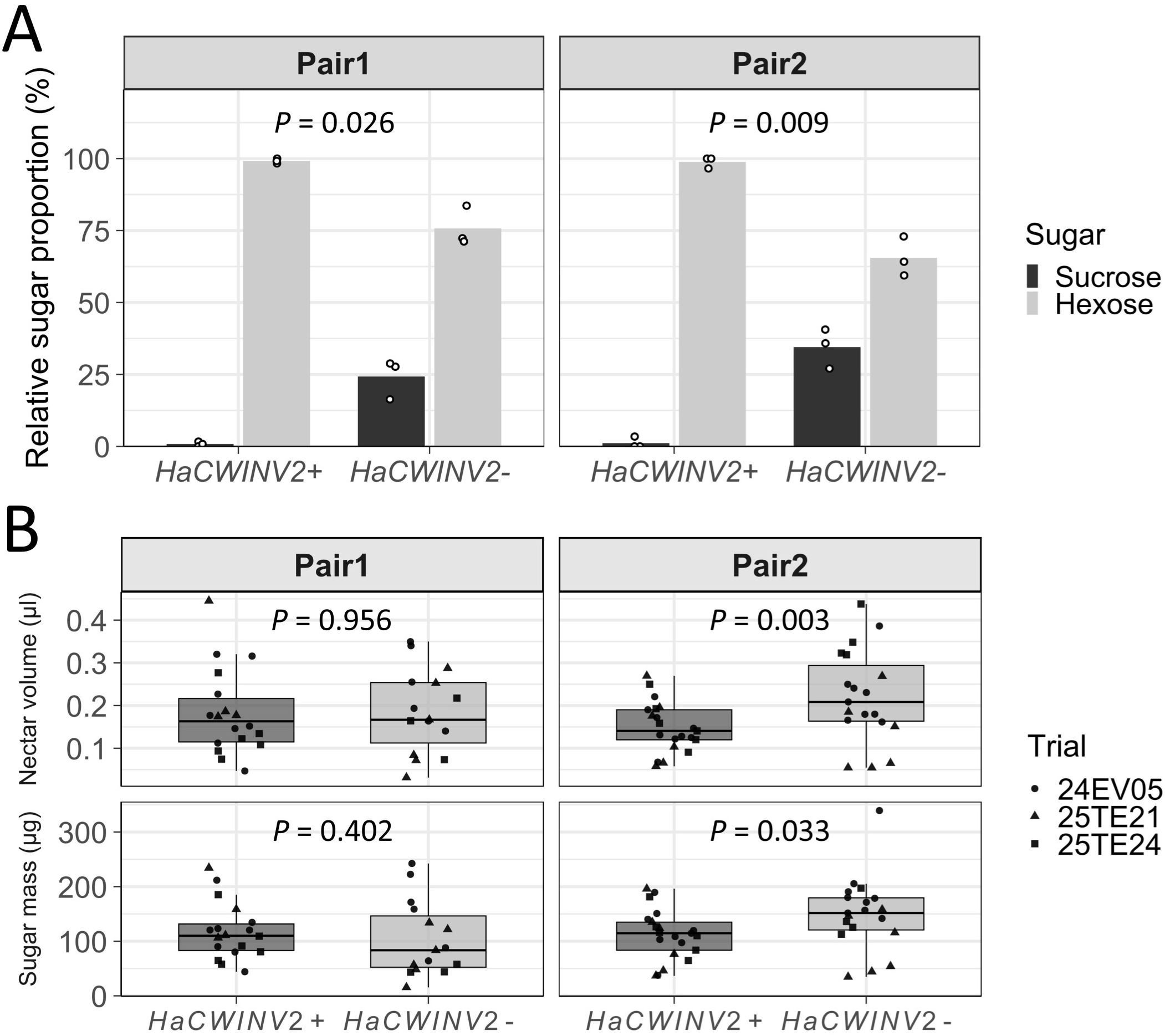
Effect of *HaCWINV2* on nectar composition, volume, and floret size in two pairs of Near Isogenic sunflower Lines. A. Mean relative proportions (%) of sucrose and hexose (glucose + fructose) in floral nectar sampled from two pairs of near isogenic lines with contrasting alleles at the *HaCWINV2* locus (n = 3 plants per allele within each NIL pair). For each plant, nectar samples from six florets were pooled. Points represent individual plants. Differences in hexose proportion between *HaCWINV2+* and *HaCWINV2−* were assessed separately within each NIL pair using Welch’s t-tests. B. Nectar volume (µl) and sugar mass (µg) per floret (plant means; n = 3–8 florets per plant). Allele effects were analysed separately within each NIL pair using linear models (ANOVA) including *HaCWINV2* allele and trial as fixed effects. P-values reported in the figure correspond to the *HaCWINV2−* vs *HaCWINV2+* contrast estimated from the fitted models using estimated marginal means (emmeans^45^). Pair 1 comprised 18 HaCWINV2+ plants and 15 *HaCWINV2−* plants; Pair 2 comprised 21 *HaCWINV2+* plants and 19 *HaCWINV2−* plants. Circles, triangles and squares denote three independent field trials conducted in 2024 and 2025.

### Genetic background, but not *HaCWINV2*, affects nectar volume, sugar mass, and floral traits

We wondered whether the alleles of *HaCWINV2* also affected nectar volume and sugar content as proposed by Minami et al. (2021)^11^. We measured nectar volume and total sugar mass from field-grown *HaCWINV2+* and *HaCWINV2−* plants in two NIL pairs and analysed each pair separately using estimated marginal means averaged over trials. In Pair 1, nectar volume and total sugar mass did not differ between alleles (*HaCWINV2*−: 0.178 µL per floret; *HaCWINV2+*: 0.180 µL per floret; *P* = 0.956; and 97.9 vs 115.4 µg per floret; *P* = 0.402, respectively) (Fig. 2B). In contrast, in Pair 2, *HaCWINV2−* plants produced larger volumes of nectar than *HaCWINV2+* plants (0.228 vs 0.149 µL per floret; +52.8%; *P* = 0.003). This was accompanied by higher total sugar mass (145 vs 108 µg per floret, +34.1%; *P* = 0.033) (Fig. 2B).

We further tested whether floret morphology varied according to *HaCWINV2* genotype. Using plant-level means (n = 12 plants per pair), floret morphology did not differ between HaCWINV2 genotypes in either Pair 1 (length *P* = 0.693; width *P* = 0.807) or Pair 2 (width *P* = 0.122; length 6.53 vs 6.33 mm, *P* = 0.064) (Supplementary Fig. 7). Overall, allele effects on nectar production were background-dependent and not consistently replicated across the two NIL pairs, while floret size showed no significant allele-dependent difference.

### Hexose-rich nectar correlates with more pollinator visits

To determine whether differences in sugar composition affected pollinator preferences, we deployed field cameras on 63 individual plants across four independent field trials to monitor insect visitation. Images were collected every 5 minutes and analysed using an ad-hoc deep-learning model to estimate the number and identity of insects visiting *HaCWINV2+* and *HaCWINV2-* flowers. Images recorded outside the daytime period (06:00–21:59) and those corresponding to late flowering were subsequently filtered out. The final dataset comprised 114,759 photographs (152 h of observation per plant on average) (Supplementary Table 8), in which we detected 28,175 bees and 2,034 bumblebees. These data were summarized into 9,599 plant × hour observations by aggregating detection events across all images taken from a given plant within each date–hour time bin. We analysed bee and bumblebee counts using generalized linear mixed models fitted separately for each NIL pair. In each model, the number of insect visits per picture was explained by the *HaCWINV2* allele while accounting for variation in weather conditions (light, temperature, humidity and wind), for repeated measurements on the same plants, and field trial. The number of images per condition was included to account for uneven sample sizes. Analysis of residuals supported an adequate fit for all negative-binomial GLMMs (Supplementary Table 4). Mixed-model R² values differed between pollinator groups: for bees, fixed effects explained 30.6% of the variance (marginal R²), increasing to 43.6% when random intercepts were included (conditional R²), whereas for bumblebees the variance explained increased from 3.6% to 18.6%.

*HaCWINV2+* flowers received 33.2% more bee visits overall (*P* = 0.0006) (Fig. 3). Estimated marginal means averaged across pairs were 0.289 and 0.217 bees per picture for *HaCWINV2+* and *HaCWINV2−* plants, respectively. The difference in visitation was consistent across both NIL pairs, with *HaCWINV2+* plants of NIL pair 1 receiving 53.6% more bee visits (0.344 vs 0.224 bees per picture; *P* = 0.0005). *HaCWINV2+* plants of NIL pair 2 also received on average 14.7% more visits (0.242 vs 0.211 bees per picture), although in isolation the effect was not statistically significant (*P* = 0.216). In contrast, there was no evidence that bumblebee visitation differed between alleles either overall (*P* = 0.758) or within each pair (Fig. 3).

**Figure 3.**
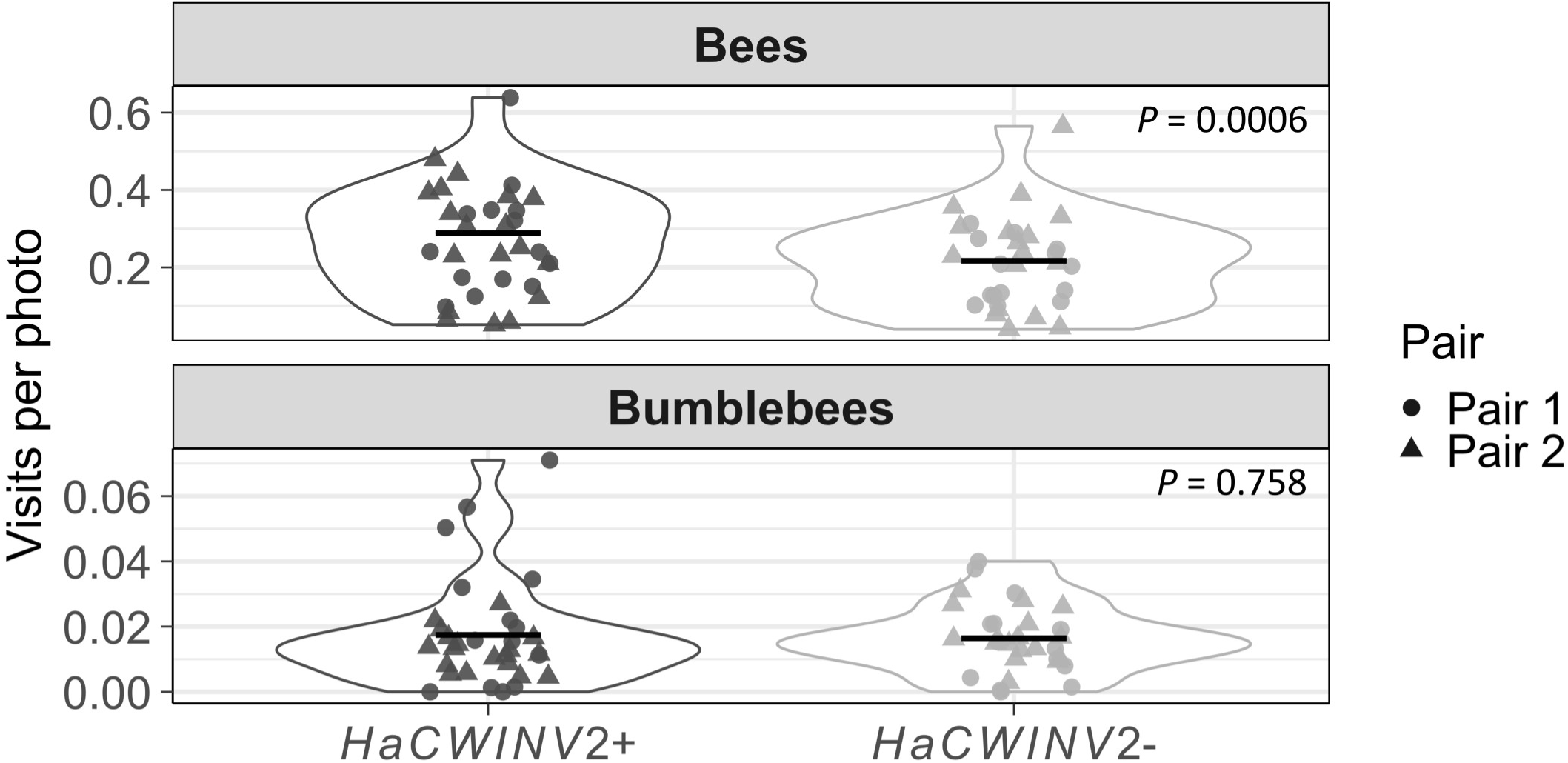
Raw pollinator counts per sunflower line differing at the *HaCWINV2* locus. Mean visitation rates per photograph for bees and bumblebees across field trials are shown for the two near-isogenic sunflower lines differing at the *HaCWINV2* locus. For each plant, detections were summed across all photographs and divided by the total number of images to obtain a plant-level mean number of visits per photo. Violin plots show the distribution of these plant-level values for each allele, and overlaid symbols represent individual plants. Black horizontal bars indicate model-estimated mean visitation rates per photograph (estimated marginal means) derived from the negative-binomial generalized linear mixed models described in the main text. Colours indicate allele (*HaCWINV2+* dark grey, *HaCWINV2−* light grey), and symbol shapes indicate NIL pair. Sample sizes were: Pair 1, n = 14 plants per allele; Pair 2, n = 18 *HaCWINV2+* plants and n = 17 *HaCWINV2−* plants.

### The *HaCWINV2* allele impacts nectar microbial diversity

We wondered how the difference in nectar sugar composition between *HaCWINV2+* and *HaCWINV2-* plants impacts pollinator visits and if this could be due to nectar microbiota as nectar volume and sugar concentration could be ruled out. To test this, we collected nectar samples from field-grown *HaCWINV2+* and *HaCWINV2-* plants belonging to NIL pairs 1 and 2. To ensure that sufficient quantities of nectar were collected, and to limit inoculation from asymmetrical pollinator visits between lines, we enclosed flower heads into nets before anthesis. We profiled bacterial and fungal communities using long-read amplicon sequencing of the bacterial 16S rRNA gene and fungal internal transcribed spacer (ITS).

The 47 16S amplicon samples were represented on average by 27,644 reads, which clustered into 13 Operational Taxonomic Units (OTUs) per sample. The dominant OTU (90c39638c0a1a1b6a2ca253c711811238359c694) accounted on average for 90% of sequenced reads in 2023 and 97% in 2024 (Supplementary Fig. 8, Supplementary Data). The representative sequence of this OTU displayed 100% identity to the *H. annuus* chloroplast 16S rRNA gene sequence. Removal of OTUs that were potential contaminants found in control samples, removed the totality of the data from all but 4 samples (Supplementary Table 9). This high proportion of plastid DNA detected in our samples might stem from low bacterial densities, higher than expected levels of plastid contamination of nectar samples, or PCR artefacts. To distinguish between these possibilities, we analysed serial dilutions of a known mixture of microbial DNA into field-collected sunflower nectar. We were able to detect sequences corresponding to as few as 8 bacterial cells, indicating that competition with chloroplastic DNA did not significantly impede amplification of bacterial markers (Supplementary Fig. 9). Instead, these results indicate that bacteria were scarce in our nectar samples. To confirm this, we collected 24 nectar samples in the field in 2024, and spread them on general bacteriological medium. Of the 24 samples tested, 6 showed no evidence of microbial growth (0 colonies after 48 hours of incubation), and the remaining 18 displayed on average 4.78 colonies per sample (Supplementary Table 10). Together, these data confirm that few bacteria were present in the field-collected nectar of netted plants.

In contrast, we detected significant fungal diversity in samples analysed using fungal ITS sequences (Fig. 4A, Supplementary Fig. 10). The 47 samples contained on average 31,618 reads and 15.11 OTUs per sample (Supplementary Data). Removal of OTUs detected in controls resulted in the exclusion of 62 OTUs, yielding 38 samples with on average 13,415 sequencing reads and 10.58 OTUs per sample (Supplementary Table 9). Rarefaction analysis further confirmed that we sampled most of the taxonomic diversity (Supplementary Fig. 3).

**Figure 4:**
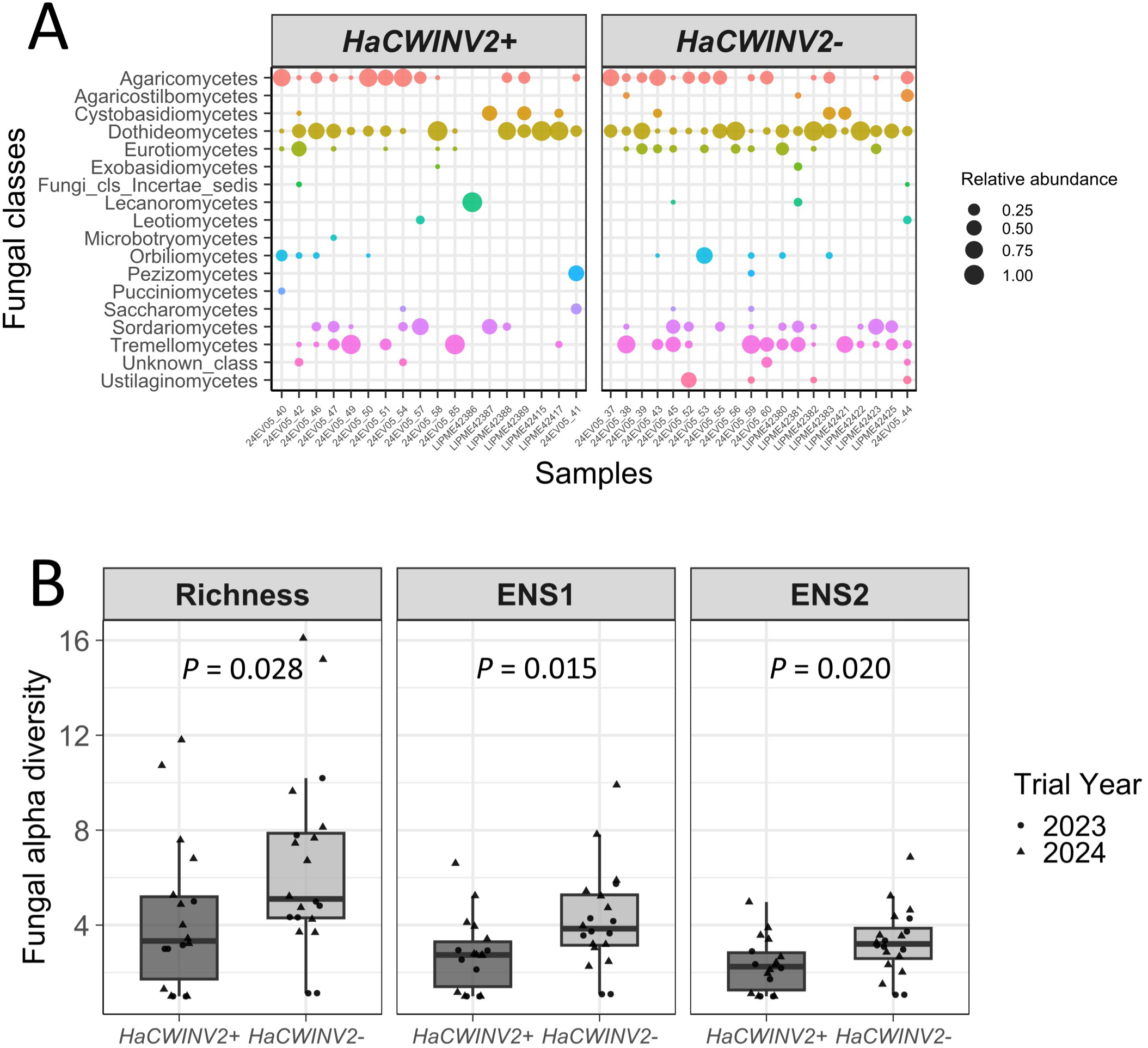
Effect of *HaCWINV2* on nectar fungal community composition and diversity. A. Relative abundance of fungal classes across individual nectar samples, stratified by *HaCWINV2* allelic class. OTUs are grouped at the class level (unclassified taxa shown as “Unknown class”). Colours indicate fungal class. Bubble size is proportional to relative abundance within each sample. Classes are ordered by prevalence across samples. Panels correspond to the two *HaCWINV2* allelic classes (*HaCWINV2+* and *HaCWINV2−*). B. OTU-level fungal alpha-diversity metrics in nectar samples from *HaCWINV2+* and *HaCWINV2−* lines. Points represent individual samples; circles and triangles indicate the field trial year. For each metric, data were analysed using ANOVA including *HaCWINV2* allele, NIL pair, their interaction (allele × pair), and trial year as fixed effects. Richness, ENS1 and ENS2 correspond to Hill numbers for richness, effective number of species calculated based on Shannon entropy, and effective number of species cal-culated based on inverse Simpson index, respectively. *P*-values reported in the figure correspond to the *HaCWINV2−* vs *HaCWINV2+* contrast estimated from statistical models using estimated marginal means (emmeans^45^). Richness was square-root transformed prior to analysis. Pair 1, n = 10 plants per allele; Pair 2, n = 8 *HaCWINV2+* plants and n = 10 *HaCWINV2−* plants.

Fungal diversity was higher in nectar of flowers carrying the *HaCWINV2−* allele than in *HaCWINV2+* flowers across both NIL pairs. When controlling for NIL pair, trial year, and the allele × pair interaction, post hoc estimated marginal mean contrasts indicated significantly higher diversity in *HaCWINV2−* nectar for all three metrics: richness (*P* = 0.028), ENS1 (*P* = 0.015), and ENS2 (*P* = 0.020) (Fig. 4B). Trial year significantly affected richness (*P* = 0.0299), showed a marginal effect on ENS1 (*P* = 0.0665), and had no detectable effect on ENS2 (*P* = 0.1233). NIL pair and the allele × pair interaction were not significant for any metric (P ≥ 0.10 for NIL pair; P ≥ 0.63 for the interaction), indicating that the *HaCWINV2* allelic effect on fungal diversity was consistent across NIL pairs and trial years.

The *HaCWINV2* allele had a small but significant effect on community structure, as detected with both Bray–Curtis distances (PERMANOVA *P* = 0.0459, R^2^ = 0.032) (Supplementary Fig. 11) and Jaccard distances (*P* = 0.0384, R^2^ = 0.036) (Supplementary Fig. 12). Trial year also had a significant effect on Jaccard distances (*P* = 0.0325, R^2^ = 0.036), whereas NIL pair had no significant main effect for either metric (*P* ≥ 0.63). To further characterize this allele-associated shift in community composition, we examined OTU-level occurrence patterns and tested for differential abundance of each OTU between *HaCWINV2* alleles and for each NIL pair. We identified 31 OTUs (15.2% of 204) that were detected in at least one *HaCWINV2+* and one *HaCWINV2−* sample. The remaining OTUs were allele-specific, with 102 OTUs (50.0%) detected only in *HaCWINV2*− samples and 71 OTUs (34.8%) detected only in *HaCWINV2+* samples. Allele-specific OTUs accounted for a large fraction of sequencing reads, representing on average 79.3% of sequences in *HaCWINV2−* samples and 68.6% of sequences in *HaCWINV2+* samples. Overall, these data support a fungal community shift driven by the sugar composition of nectar and the *HaCWINV2* allele.

## Discussion

Animal-mediated pollination depends on variation in floral traits that shape visitor behaviour. Yet, establishing direct mechanistic links between genetic variation, reward chemistry and pollinator responses remains challenging, because floral traits often covary and their ecological effects are context dependent. Disentangling how specific components of floral rewards influence both visitor behaviour and associated microbial communities is therefore essential to connect genotype to ecological function in plant–pollinator interactions.

We show here that *HaCWINV2,* coding for a cell wall invertase of *H. annuus,* contributes to nectar sugar chemistry. A naturally occurring variant of *HaCWINV2*, which presents a non-synonymous substitution resulting in the replacement of a conserved arginine residue by a cysteine at position 191, is catalytically inactive when expressed in yeast. Sunflower lines homozygous for the R191C allele produce nectar with elevated sucrose levels relative to near-isogenic lines carrying the functional allele, demonstrating that *HaCWINV2* directly controls nectar sugar composition. In contrast to what has been reported in *Arabidopsis* and *Brassica*, plants carrying the catalytically impaired *HaCWINV2* allele did not consistently produce lower nectar volumes or total sugar amounts^10,11^. In Brassicaceae, hexose-rich nectar production has been linked to cell wall invertases that hydrolyse sucrose and generate a concentration gradient driving sugar influx into nectaries^11^. The absence of a strong relationship between nectar volume and sucrose proportion in sunflower suggests that nectar secretion may rely on alternative mechanisms that are independent of a sucrose-driven gradient^11^.

Flower attractiveness to pollinators is influenced by floret size and nectar quantity^5,24^. However, whether nectar sugar composition alone affects pollinator behaviour has been more difficult to address because chemical composition is often linked to other floral traits^1,11^. Our results demonstrate that near-isogenic lines that produced hexose-rich nectar attract significantly more bees than lines with identical floral traits that produced sucrose-rich nectar. Across four field trials over three years and 114,759 images, there was strong evidence that the presence of a *HaCWINV2+* allele increased bee visits overall (+33%). This increase in bee visits was observed for *HaCWINV2+* flowers of both NIL pairs, although we were only able to establish statistical significance for NIL pair 1. The less pronounced effect detected for NIL pair 2 may be due to a bias in nectar production, as the *HaCWINV2-* line of pair 2 produced more nectar (+52.8% volume) and total sugar (+34.1%). Higher nectar sugar content is an established factor positively influencing bee visits, including on sunflower^5^. However, if the total sugar mass per floret was the main driver of attractiveness, higher visitation rates would have been expected on *HaCWINV2−* plants with this genetic background, which was not observed. Together, our data clearly demonstrate a preference of bees for *HaCWINV2+* hexose-rich nectar. This preference for *HaCWINV2+* flowers likely confers a selective advantage in wild sunflower populations. Indeed, our survey of sunflower genetic diversity indicated that alleles linked to the *HaCWINV2* R191C loss-of-function substitution are rare at about 3% in wild populations, and only maintained in a heterozygous state. This indicates that the recessive loss-of-function *HaCWINV2* allele is likely counter-selected in the wild. However, catalytically-impaired alleles of *HaCWINV2* are frequent in cultivated inbred lines (34.8%), despite a measurable deleterious effect on pollinator attractiveness. This pattern may reflect a relaxation of pollinator-mediated selection under domestication. Sunflower cultivars are largely self-compatible and self-fertile, with pollinator dependency varying widely among cultivars and environments^62^. Because commercial sunflower fields are typically planted as single cultivars, pollinators may have limited opportunities to choose among genotypes differing in nectar composition. Under such non-choice conditions, the relative preference observed in experimental comparisons can exert weaker selective pressure, although this effect remains relevant in seed production systems where different parental lines are grown together. This reduces the selective cost of reduced floral attractiveness in agricultural contexts. Thus, the fixation of loss-of-function *HaCWINV2* alleles in sunflower cultivars could have occurred neutrally through the selection for other genetically linked, desirable agronomic traits. Alternatively, *HaCWINV2* may incur a trade-off with other traits that may have been selected during breeding. A plausible trade-off may involve resistance to pathogens. Indeed, cell wall invertases are involved in plant immunity and sugar-mediated defence reprogramming in *Arabidopsis*, with functional perturbations affecting defence signalling^63,64^. Another intriguing possibility may involve the selection of specific microbiota in hexose vs. sucrose-rich nectar, which are passed on to the seed, thereby influencing agriculturally-relevant traits such as seed performance and germination^65,66^.

In contrast to the clear effect of *HaCWINV2* variants on bee visits, bumblebees did not display significant preferences for plants expressing either allele. This absence of response suggests that nectar sugar composition does not universally determine pollinator behaviour, but instead contributes to a multidimensional reward landscape. Bees and bumblebees may respond differently to floral traits, including reward and accessibility traits, and such non-parallel responses are expected^7,8^. Historically, studies in sunflower reported a preference of foragers for sucrose-rich nectar^67^, implying a direct role of sugar identity in shaping attractiveness. However, nectar quality is multidimensional, and sugar composition rarely varies independently from concentration, viscosity and other chemical constituents, including amino acids and secondary metabolites^1^. Because bees generally exploit both sucrose- and hexose-rich nectars^1^, differences in visitation are unlikely to reflect an intrinsic preference for particular sugars. Instead, sugar identity may influence visitation indirectly by reshaping the physicochemical environment of nectar and the chemical cues associated with it, including through effects on microbial communities and their volatile compounds.

Nectar-associated microbes modify nectar chemistry and emit volatile compounds, thereby altering the sensory landscape perceived by pollinators^13^. Consistent with this hypothesis, we detected a significant influence of nectar sugar composition on fungal communities in pairs of near-isogenic lines differing in nectar sucrose content. Fungal communities in sucrose-rich (*HaCWINV2−*) nectar were significantly more diverse than those in hexose-rich (*HaCWINV2+*) nectar. This difference was driven by limited overlap in OTU composition between alleles and by a high proportion of allele-specific OTUs, which accounted for a substantial fraction of sequencing reads (79.3% of reads per sample in *HaCWINV2−* and 68.6% in *HaCWINV2+*). These patterns are consistent with stochastic community assembly in an ephemeral habitat such as nectar, a process likely amplified here by pollinator exclusion^15^. Sucrose-rich nectar nevertheless harboured more diverse and compositionally distinct fungal communities. These results suggest that sugar composition acts as an environmental filter. Sucrose may increase resource heterogeneity and modify osmotic conditions, thereby relaxing physiological constraints and enabling the establishment of a broader range of fungal taxa. This interpretation is consistent with experimental work in artificial nectars showing that sugar concentration, composition and nutrient availability constrain microbial community assembly^18^. Differences in microbial diversity induced by *HaCWINV2* variation may in turn modulate the reliability of microbe-derived olfactory cues, with more specialized communities potentially generating more consistent volatile signatures^13^. In parallel, higher diversity may also enhance resistance to pathogen invasion via niche saturation or competitive exclusion^68^. In open flowers, repeated pollinator visits further structure nectar microbial communities^15,16,69^. Because heads were netted in our experiments to avoid bias due to uneven insect-mediated transport of microbiota, the communities described in this study likely differ from those occurring when pollinators have free access. Nevertheless, our results provide evidence of ecological filters at the level of nectar composition, driven by plant genotype which would be expected to hold under different pollinator access regimes.

In conclusion, we identified *HaCWINV2* as a key genetic driver of plant–nectar–pollinator interactions. Natural variation at this single locus shapes nectar chemistry, with cascading effects on nectar and pollinator visitation. Whether microbiota, perhaps through the production of volatile compounds, directly contributes to the preference of bees for hexose-rich sunflower nectar, remains an open question.

## ACKNOWLEDGMENTS

This work is part of HELEX project funded by the European Union’s Horizon Europe Research and Innovation Actions programme under grant agreement N°101081974. This research used the PHENOME-EMPHASIS facility Phenotoul-Polliphen (Phenome-ANR-11-INBS-0012, https://doi.org/10.15454/1.5483266728434124E12) and was part of the French Laboratory of Excellence project “TULIP“ (ANR-10-LABX-41; ANR-11-IDEX-0002-02). This work was funded by the Plant2Pro Carnot Institute in the frame of its 2022 call for projects. Plant2Pro is supported by ANR (agreement #22-CARN-024-01 – 2021). This work was supported by the International Consortium of Sunflower Genomics and partially funded by the Promosol project Heliopollen. G.T. acknowledges support from the INRAE BAP department. Cultivated lines resequencing of the SEAM population was funded by the Agence Nationale de Recherche, programme France2030’s project AgroDiv (ANR-22-PEAE-0005) and wild *H. annuus* resequencing was funded from Promosol project Heliawild. We are grateful to the Genotoul bioinformatics platform Toulouse Occitanie (Bioinfo Genotoul, https://doi.org/10.15454/1.5572369328961167E12) for providing help and/or computing and/or storage resources. We are grateful to Baptiste Mayjonade for help and advice on ONT Nanopore sequencing. We are grateful to Mireille Haon and Sacha Grisel from the 3PE platform (Marseille, France) for their help with yeast cultures, as well as Nico Callewaert (VIB-UGent) for kindly sharing the OpenPichia strains and plasmids.

## AUTHOR CONTRIBUTIONS

G.T., N.L., and A.C. conceived the study. N.P., A.R. and E.L. cloned recombinant genes into *K. phaffii*, assessed protein expression and enzymatic activity. N.P. developed the NIL populations and performed PACE and Axiom genotyping analyses. N.L. analysed RNA-seq data. N.P. and A.L. extracted nucleic acids used in sequencing experiments and genotyping. J.B.-S. built and tested the image analysis software. G.T., N.L., N.B., and O.C. annotated photographs used in training the image analysis model. S.C. and J.G. contributed genome sequence analysis. M.M. performed variant calling analysis of sunflower genome sequencing data. G.T., M.-P.J., and E.L. performed sugar enzymatic assays. M.-C.B. managed the field trials. G.T. sampled nectar in the field and quantified nectar traits. O.C. measured floret morphological traits. G.T., S.M. and N.P. extracted nucleic acids from nectar samples, performed PCR and sequencing. G.T. and A.C. analysed amplicon sequencing data. G.T. performed statistical analyses. G.T., A.C., and N.L. wrote the manuscript with contributions from all authors. A.C. and N.L. contributed equally to the supervision of the work.

## DATA STATEMENT

All sequence data generated during the study available in a public repository, accession numbers are provided in the main text.

Raw data on nectar and floral traits, pollinator visits, sequence variants at the *HaCWINV2* locus, and OTU tables are available on the INRAE institutional repository at https://doi.org/10.57745/PRMSCV

## REFERENCES

1. Nicolson, S. W. Sweet solutions: nectar chemistry and quality. Philos. Trans. R. Soc. B Biol. Sci. 377, 20210163 (2022).

2. Roy, R., Schmitt, A. J., Thomas, J. B. & Carter, C. J. Review: Nectar biology: From molecules to ecosystems. Plant Sci. 262, 148–164 (2017).

3. Mallinger, R. & Prasifka, J. Benefits of Insect Pollination to Confection Sunflowers Differ Across Plant Genotypes. Crop Sci. 57, 3264–3272 (2017).

4. Greenleaf, S. S. & Kremen, C. Wild bees enhance honey bees’ pollination of hybrid sunflower. Proc. Natl. Acad. Sci. 103, 13890–13895 (2006).

5. Mallinger, R. E. & Prasifka, J. R. Bee visitation rates to cultivated sunflowers increase with the amount and accessibility of nectar sugars. J. Appl. Entomol. 141, 561–573 (2017).

6. Bartomeus, I. et al. Contribution of insect pollinators to crop yield and quality varies with agricultural intensification. PeerJ 2, e328 (2014).

7. Mu, J. et al. Honey bees and bumble bees react differently to nitrogen-induced increases in floral resources. Environ. Entomol. 53, 1111–1119 (2024).

8. Verweij, F., Biesmeijer, K. & Klumpers, S. Differences in visitation of honeybees and bumblebees to ornamental plant varieties can be explained by floral traits. J. Pollinat. Ecol. 38, 36–57 (2025).

9. Lin, I. W. et al. Nectar secretion requires sucrose phosphate synthases and the sugar transporter SWEET9. Nature 508, 546–549 (2014).

10. Ruhlmann, J. M., Kram, B. W. & Carter, C. J. CELL WALL INVERTASE 4 is required for nectar production in Arabidopsis. J. Exp. Bot. 61, 395–404 (2010).

11. Minami, A., Kang, X. & Carter, C. J. A cell wall invertase controls nectar volume and sugar composition. Plant J. 107, 1016–1028 (2021).

12. Krömer, T., Kessler, M., Lohaus, G. & Schmidt-Lebuhn, A. N. Nectar sugar composition and concentration in relation to pollination syndromes in Bromeliaceae. Plant Biol. 10, 502–511 (2008).

13. Rering, C. C., Beck, J. J., Hall, G. W., McCartney, M. M. & Vannette, R. L. Nectar-inhabiting microorganisms influence nectar volatile composition and attractiveness to a generalist pollinator. New Phytol. 220, 750–759 (2018).

14. Martin, V. N., Schaeffer, R. N. & Fukami, T. Potential effects of nectar microbes on pollinator health. Philos. Trans. R. Soc. B Biol. Sci. 377, 20210155 (2022).

15. Vannette, R. L. & Fukami, T. Dispersal enhances beta diversity in nectar microbes. Ecol. Lett. 20, 901–910 (2017).

16. Morris, M. M., Frixione, N. J., Burkert, A. C., Dinsdale, E. A. & Vannette, R. L. Microbial abundance, composition, and function in nectar are shaped by flower visitor identity. FEMS Microbiol. Ecol. 96, fiaa003 (2020).

17. Mueller, T. G., Francis, J. S. & Vannette, R. L. Nectar compounds impact bacterial and fungal growth and shift community dynamics in a nectar analog. Environ. Microbiol. Rep. 15, 170–180 (2023).

18. Pozo, M. I. et al. Species coexistence in simple microbial communities: unravelling the phenotypic landscape of co-occurring Metschnikowia species in floral nectar. Environ. Microbiol. 18, 1850–1862 (2016).

19. Verhaest, M. et al. X-ray diffraction structure of a cell-wall invertase from Arabidopsis thaliana. Acta Crystallogr. D Biol. Crystallogr. 62, 1555–1563 (2006).

20. Lammens, W. et al. Structural insights into glycoside hydrolase family 32 and 68 enzymes: functional implications. J. Exp. Bot. 60, 727–740 (2009).

21. Reddy, A. & Maley, F. Studies on identifying the catalytic role of Glu-204 in the active site of yeast invertase. J. Biol. Chem. 271, 13953–13957 (1996).

22. Nagem, R. a. P., et al. Crystal structure of exo-inulinase from Aspergillus awamori: the enzyme fold and structural determinants of substrate recognition. J. Mol. Biol. 344, 471–480 (2004).

23. Meng, G. & Fütterer, K. Structural framework of fructosyl transfer in Bacillus subtilis levansucrase. Nat. Struct. Biol. 10, 935–941 (2003).

24. Prasifka, J. R. et al. Using Nectar-Related Traits to Enhance Crop-Pollinator Interactions. Front. Plant Sci. 9, 812 (2018).

25. Barstow, A. C., Prasifka, J. R., Attia, Z., Kane, N. C. & Hulke, B. S. Genetic mapping of a pollinator preference trait: Nectar volume in sunflower (Helianthus annuus L.). Front. Plant Sci. 13, (2022).

26. Barstow, A. C. et al. Variant filters using segregation information improve mapping of nectar-production genes in sunflower (Helianthus annuus L.). Plant Genome 18, e70042 (2025).

27. Cruden, R. W., Hermann, S. & Peterson, S. Patterns of nectar production and plant-pollinator coevolution. in The biology of nectaries. Pp. 80–125 (Columbia University Press, New York, 1983).

28. Schneider, C. A., Rasband, W. S. & Eliceiri, K. W. NIH Image to ImageJ: 25 years of image analysis. Nat. Methods 9, 671–675 (2012).

29. Gosseau, F. et al. Heliaphen, an outdoor high-throughput phenotyping platform designed to integrate genetics and crop modeling. 362715 Preprint at 10.1101/362715 (2018).

30. Patro, R., Duggal, G., Love, M. I., Irizarry, R. A. & Kingsford, C. Salmon provides fast and bias-aware quantification of transcript expression. Nat. Methods 14, 417–419 (2017).

31. Huang, K., et al. The genomics of linkage drag in inbred lines of sunflower. Proc. Natl. Acad. Sci. 120, e2205783119 (2023).

32. Badouin, H. et al. The sunflower genome provides insights into oil metabolism, flowering and Asterid evolution. Nature 546, 148–152 (2017).

33. Li, H. et al. The Sequence Alignment/Map format and SAMtools. Bioinformatics 25, 2078–2079 (2009).

34. Paradis, E. & Schliep, K. ape 5.0: an environment for modern phylogenetics and evolutionary analyses in R. Bioinformatics 35, 526–528 (2019).

35. Katoh, K., Rozewicki, J. & Yamada, K. D. MAFFT online service: multiple sequence alignment, interactive sequence choice and visualization. Brief. Bioinform. 20, 1160–1166 (2019).

36. Claes, K. et al. OPENPichia: licence-free Komagataella phaffii chassis strains and toolkit for protein expression. Nat. Microbiol. 9, 864–876 (2024).

37. Wu, S. & Letchworth, G. J. High efficiency transformation by electroporation of Pichia pastoris pretreated with lithium acetate and dithiothreitol. BioTechniques 36, 152–154 (2004).

38. nanoporetech/dorado. Oxford Nanopore Technologies (2026).

39. De Coster, W. & Rademakers, R. NanoPack2: population-scale evaluation of long-read sequencing data. Bioinformatics 39, btad311 (2023).

40. Zheng, Z. et al. Symphonizing pileup and full-alignment for deep learning-based long-read variant calling. Nat. Comput. Sci. 2, 797–803 (2022).

41. You, F. M. et al. BatchPrimer3: A high throughput web application for PCR and sequencing primer design. BMC Bioinformatics 9, 253 (2008).

42. Dellaporta, S. L., Wood, J. & Hicks, J. B. A plant DNA minipreparation: Version II. Plant Mol. Biol. Report. 1, 19–21 (1983).

43. Milne, I. et al. Flapjack--graphical genotype visualization. Bioinforma. Oxf. Engl. 26, 3133–3134 (2010).

44. Shapiro, S. S. & Wilk, M. B. An analysis of variance test for normality (complete samples)†. Biometrika 52, 591–611 (1965).

45. Estimated marginal means (aka Least-squares means) — emmeans-package. https://rvlenth.github.io/emmeans/reference/emmeans-package.html.

46. Welch, B. L. The generalisation of student’s problems when several different population variances are involved. Biometrika 34, 28–35 (1947).

47. Brooks, M. E. et al. glmmTMB Balances Speed and Flexibility Among Packages for Zero-inflated Generalized Linear Mixed Modeling. R J. 9, 378–400 (2017).

48. Vincze, C., Leelőssy, Á., Zajácz, E. & Mészáros, R. A review of short-term weather impacts on honey production. Int. J. Biometeorol. 69, 303–317 (2025).

49. Hartig, F., Lohse, L. & leite, M. de S. DHARMa: Residual Diagnostics for Hierarchical (Multi-Level / Mixed) Regression Models. (2024).

50. Martin, M. Cutadapt removes adapter sequences from high-throughput sequencing reads. EMBnet.journal 17, 10–12 (2011).

51. Rognes, T., Flouri, T., Nichols, B., Quince, C. & Mahé, F. VSEARCH: a versatile open source tool for metagenomics. PeerJ 4, e2584 (2016).

52. Bolyen, E. et al. Reproducible, interactive, scalable and extensible microbiome data science using QIIME 2. Nat. Biotechnol. 37, 852–857 (2019).

53. Yilmaz, P. et al. The SILVA and “All-species Living Tree Project (LTP)” taxonomic frameworks. Nucleic Acids Res. 42, D643–D648 (2014).

54. Abarenkov, K. et al. UNITE general FASTA release for Fungi. UNITE Community 10.15156/BIO/2938067 (2023).

55. McMurdie, P. J. & Holmes, S. phyloseq: An R Package for Reproducible Interactive Analysis and Graphics of Microbiome Census Data. PLOS ONE 8, e61217 (2013).

56. Oksanen, J. et al. vegan: Community Ecology Package. (2025).

57. Hsieh, T. C., Ma, K. H. & Chao, A. iNEXT: an R package for rarefaction and extrapolation of species diversity (Hill numbers). Methods Ecol. Evol. 7, 1451–1456 (2016).

58. Chao, A. et al. Rarefaction and extrapolation with Hill numbers: a framework for sampling and estimation in species diversity studies. Ecol. Monogr. 84, 45–67 (2014).

59. Fox, J., et al. car: Companion to Applied Regression. (2026).

60. Lüdecke, D., Ben-Shachar, M. S., Patil, I., Waggoner, P. & Makowski, D. performance: An R Package for Assessment, Comparison and Testing of Statistical Models. J. Open Source Softw. 6, 3139 (2021).

61. Mattanovich, D. et al. Genome, secretome and glucose transport highlight unique features of the protein production host Pichia pastoris. Microb. Cell Factories 8, 29 (2009).

62. Chabert, S. et al. Importance of maternal resources in pollen limitation studies with pollinator gradients: A case study with sunflower. Agric. Ecosyst. Environ. 330, 107887 (2022).

63. Proels, R. K. & Hückelhoven, R. Cell-wall invertases, key enzymes in the modulation of plant metabolism during defence responses. Mol. Plant Pathol. 15, 858–864 (2014).

64. Tauzin, A. S. & Giardina, T. Sucrose and invertases, a part of the plant defense response to the biotic stresses. Front. Plant Sci. 5, (2014).

65. Prado, A., Marolleau, B., Vaissière, B. E., Barret, M. & Torres-Cortes, G. Insect pollination: an ecological process involved in the assembly of the seed microbiota. Sci. Rep. 10, 3575 (2020).

66. Matilla, A. J. Exploring the Bacterial Microbiota of Seeds. Microb. Biotechnol. 18, e70230 (2025).

67. Pham-Delegue, M. H. et al. Chemicals involved in honeybee-sunflower relationship. J. Chem. Ecol. 16, 3053–3065 (1990).

68. Farkas, Á., Mihalik, E., Dorgai, L. & Bubán, T. Floral traits affecting fire blight infection and management. Trees 26, 47–66 (2012).

69. Belisle, M., Peay, K. G. & Fukami, T. Flowers as islands: spatial distribution of nectar-inhabiting microfungi among plants of Mimulus aurantiacus, a hummingbird-pollinated shrub. Microb. Ecol. 63, 711–718 (2012).

