## supplementary figure for "A plant single nucleotide polymorphism impacts nectar sugar composition, microbial diversity and pollinator visits"

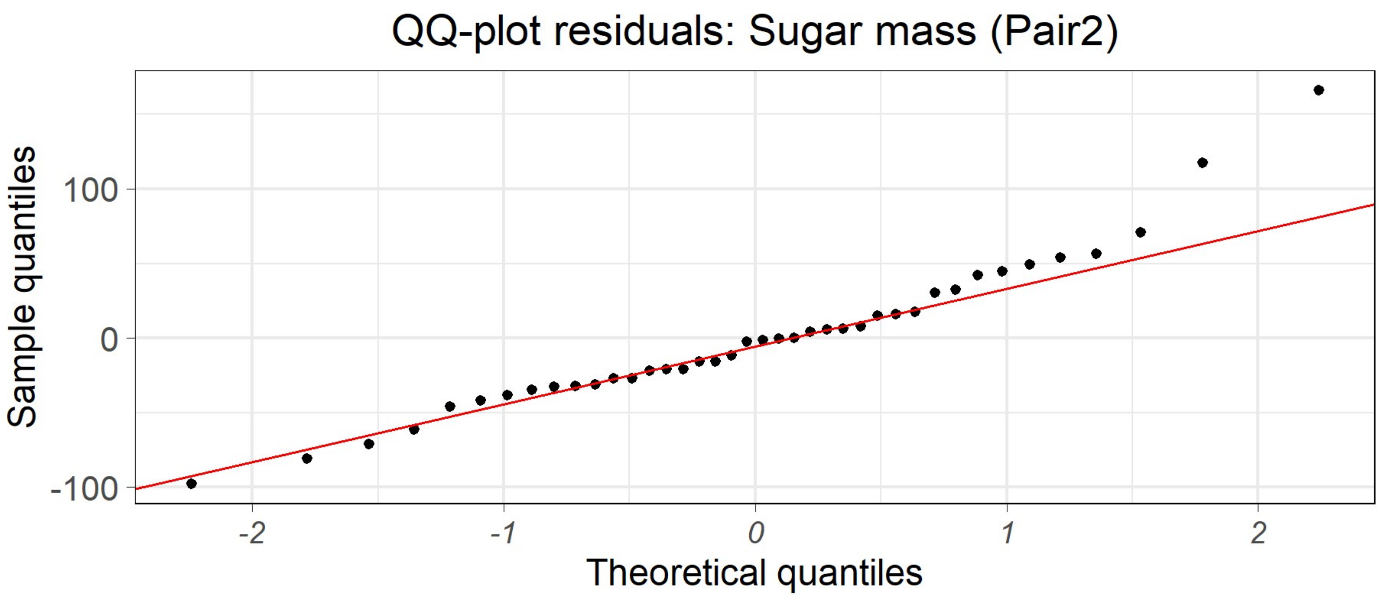


**Supplementary Figure 1. Q–Q plot of residuals from the ANOVA model fitted to plant-level mean sugar mass (µg per floret) in NIL Pair 2, including *HaCWINV2* allele and Trial as fixed effects.** Points represent standardized model residuals plotted against theoretical normal quantiles; the red line indicates the expected distribution under normality. A slight deviation is visible in the upper tail, consistent with the Shapiro–Wilk test ($P = 0.047$), but no marked departure from normality is observed.

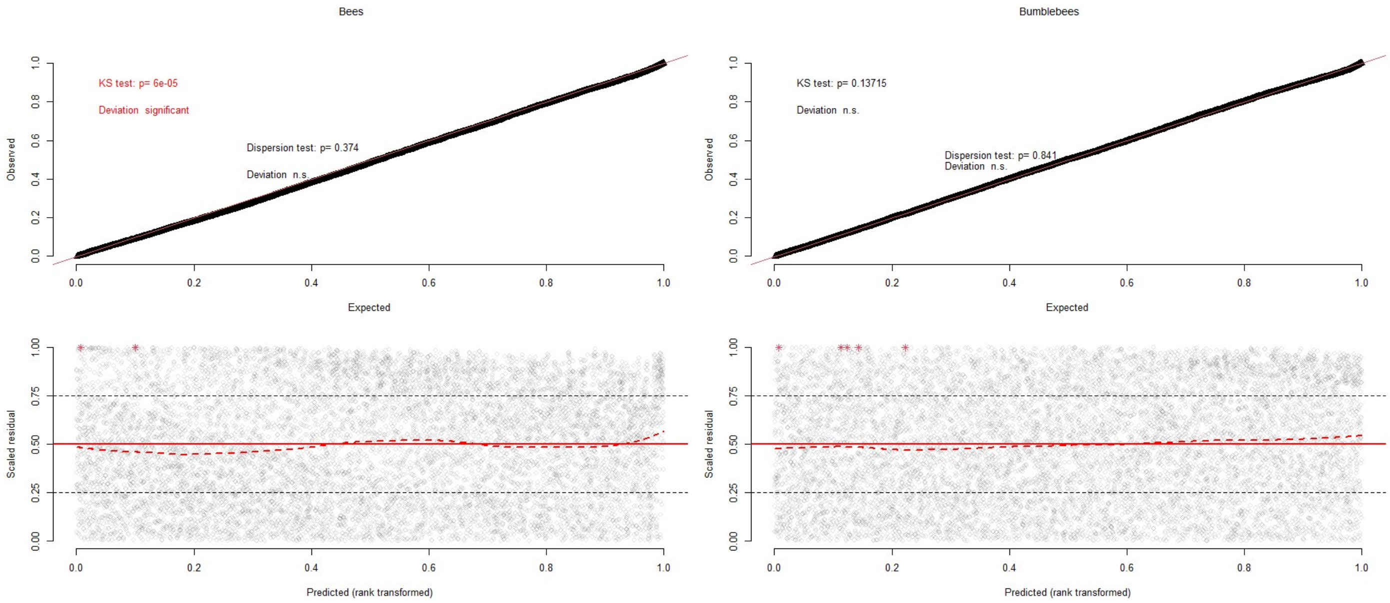


**Supplementary Figure 2. DHARMa residual diagnostics for negative binomial GLMMs of pollinator visitation.** Standardized simulated residuals were generated with DHARMa using 10,000 simulations for the two negative binomial mixed-effects models fitted to bee and bumblebee visitation data. The top row shows QQ-uniformity plots comparing scaled residuals with the expected uniform distribution. The bottom row shows scaled residuals plotted against model-predicted values. Together, these graphical diagnostics complement the formal DHARMa tests for dispersion, zero inflation, uniformity, and outliers reported in Supplementary Table 4.

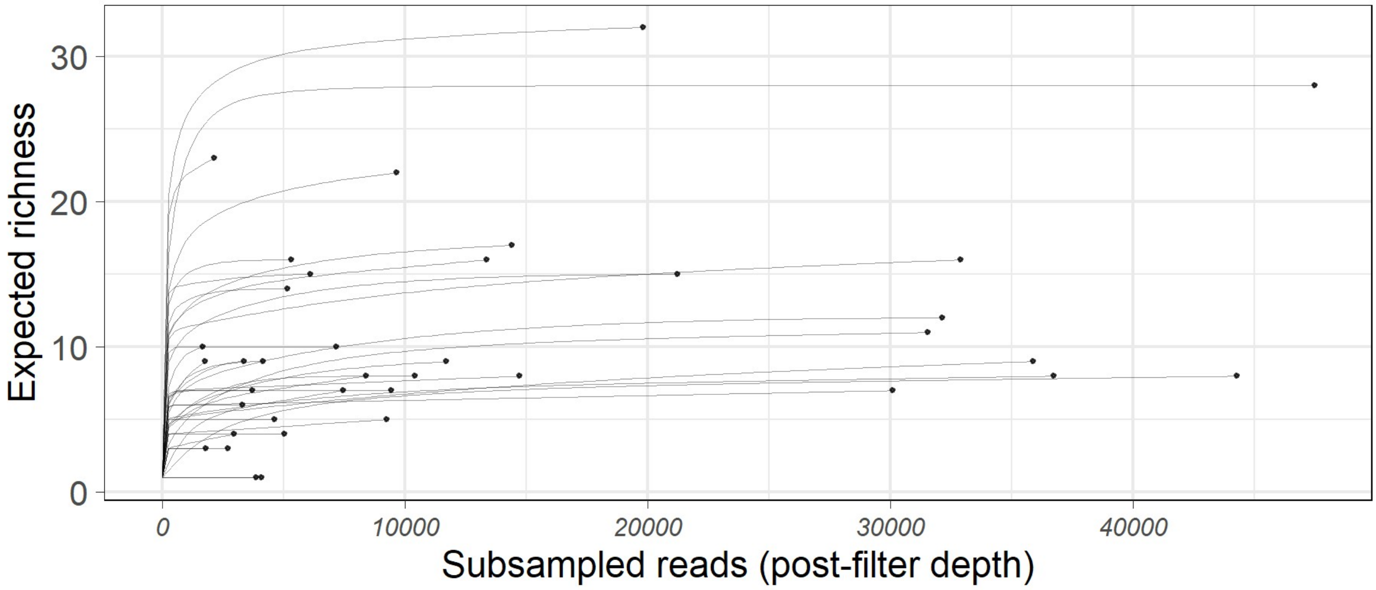


**Supplementary Figure 3. Rarefaction curves of nectar fungal communities.**
Rarefaction curves show the expected OTU richness as a function of sequencing depth for each nectar sample after contaminant removal. OTUs were first screened using negative controls (empty capillaries processed alongside samples and reagent controls) and any OTU reaching >5% relative abundance in at least one negative-control library was considered a contaminant and removed from the entire dataset. OTUs assigned to common contaminants *Malasseziomycetes* were removed. Sample sizes per NIL pair were: Pair 1, n = 10 plants per allele; Pair 2, n = 8 *HaCWINV2*+ plants and n = 10 *HaCWINV2*- plants.

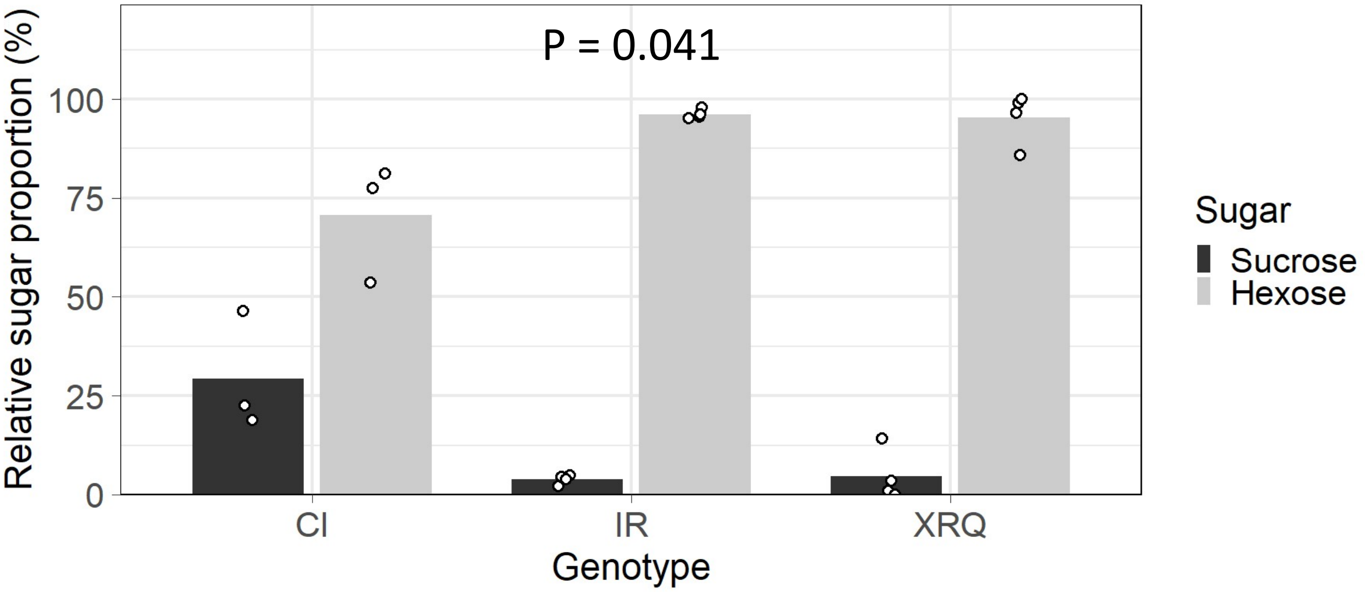


**Supplementary Figure 4. Sugar composition of nectar in CI, IR and XRQ genotypes.** Mean relative proportions of sucrose and hexoses (glucose + fructose) in nectar samples of sunflower genotypes XRQ, CI and IR. Bars indicate the mean percentage of each sugar class, and white points represent individual biological samples (one plant). For each plant, nectar from six florets was pooled, and sugar concentrations were measured using 4–5 technical replicates, averaged to obtain a single plant-level estimate. Differences among genotypes were tested using a Kruskal–Wallis test on plant-level hexose proportions.

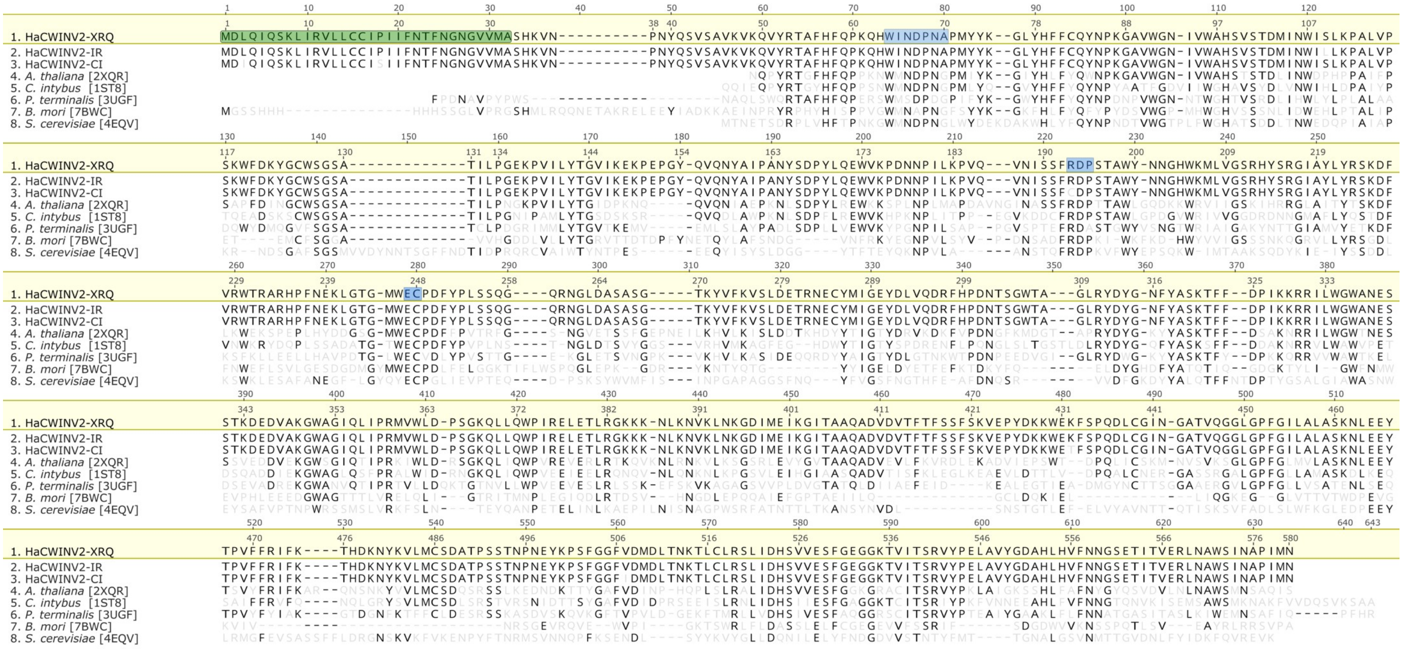

**Supplementary Figure 5. Protein sequence alignment of HaCWINV2 to experimentally-characterized sucrose invertases.** HaCWINV2 of *H. annuus* lines XRQ, CI and IR amino acid sequences were downloaded from heliagene.org (see main text for details). Protein sequences of eukaryotic representatives of the GH32 family were downloaded from RSCB PDB using the accession numbers given between brackets. *A. thaliana* [2XQR] = AtCWINV4 ; *C. intybus* [1ST8] : 1-exohydrolase IIa from *Cichorium intybus*; *P. terminalis* [3UGF] = 6-SST/6-SFT from *Pachysandra terminalis*; *B. mori* [7BWC] = *Bombyx mori* GH32 beta-fructofuranosidase BmSUC1; *S. cerevisiae* [4EQV] = *Saccharomyces cerevisiae* invertase. Sequences were aligned using MAFFT v7 in “auto” mode with default settings. Highlighted regions on the HaCWINV2-XRQ sequence denote a putative secretion signal (green), and three conserved motifs involved in catalysis or substrate binding (blue).

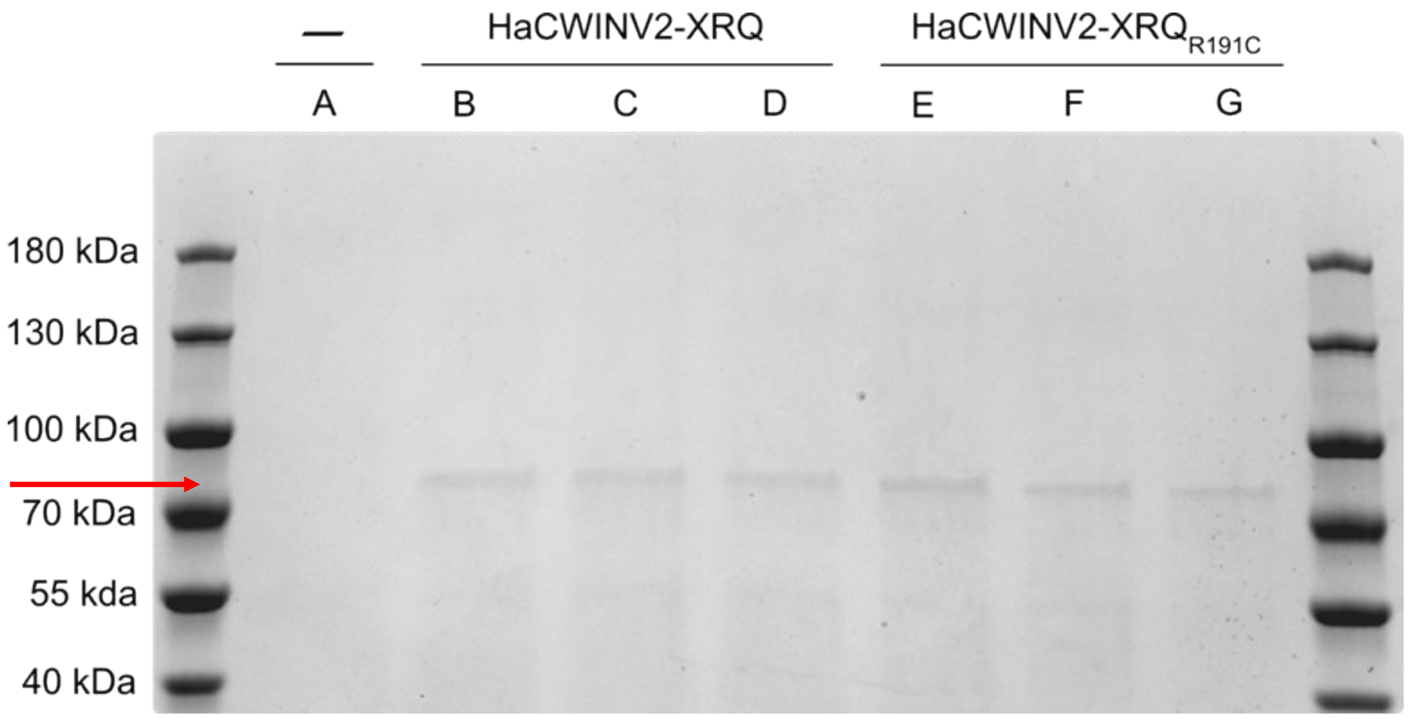


**Supplementary Figure 6. Recombinant HaCWINV2 production in K. phaffii.** Culture supernatants of K. phaffii transformants expressing HaCWINV2-XRQ or HaCWINV2-XRQR191C were analysed by SDS–PAGE. A subset of three clones per construct is shown. The expected recombinant protein band is at approximately 72.9 kDa (red arrow). “–” corresponds to the supernatants of the parental strain. Molecular-weight markers (kDa) are indicated.

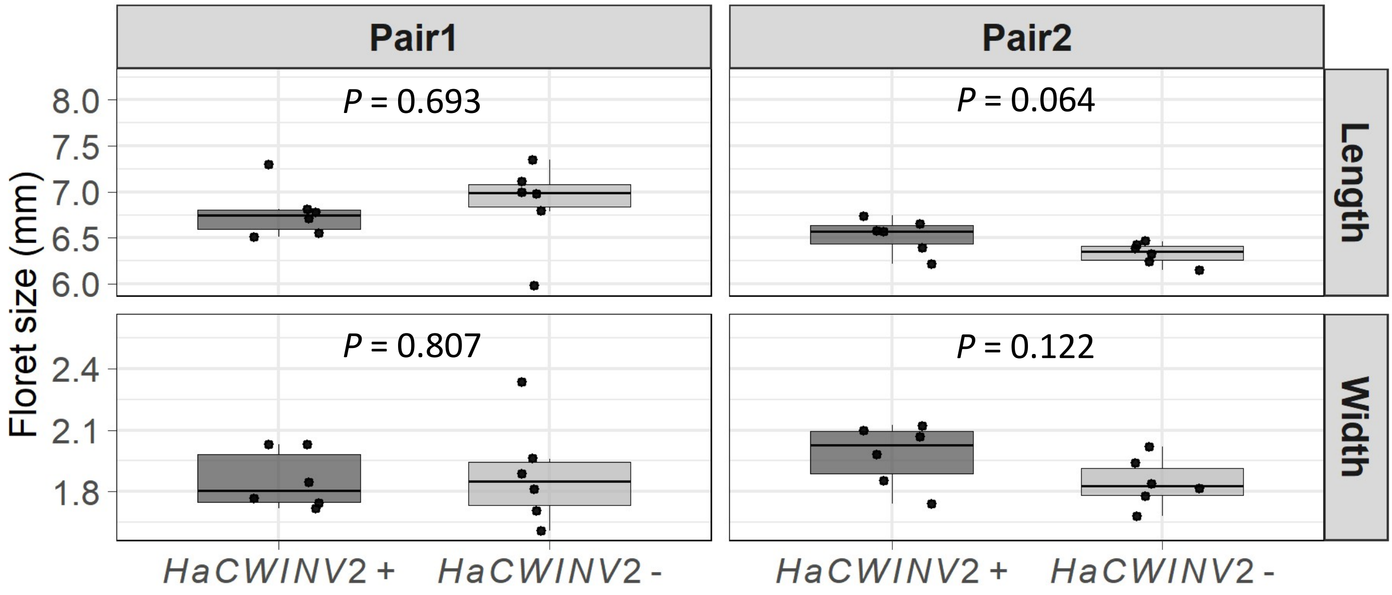


**Supplementary Figure 7. Floret length and width (mm) measured in the 24EV05 field trial and averaged at the plant level.** Boxplots show the distribution of plant means for each HaCWINV2 allele within each NIL pair. Pairwise comparisons were performed using two-sided Welch t-tests. Sample sizes were n = 6 plants per line in each NIL pair.

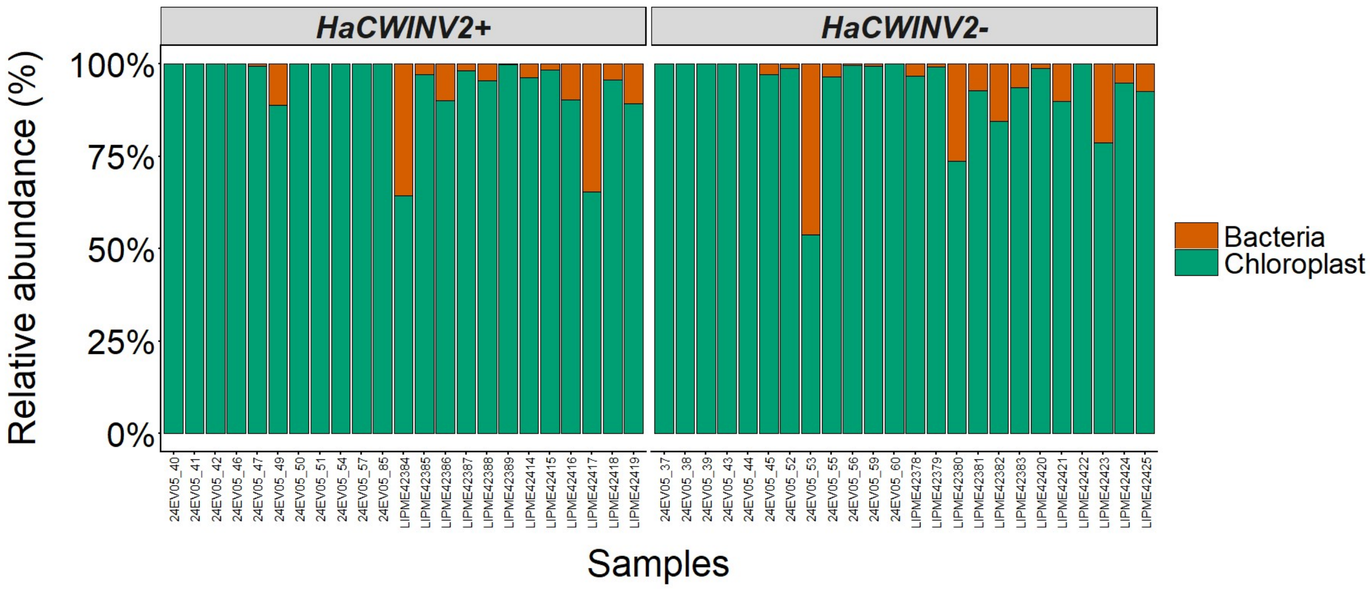


**Supplementary Figure 8. Relative abundance of chloroplast versus non-chloroplast reads per sample, stratified by *HaCWINV2* allelic class.**

Stacked barplots show the relative abundance (% of total reads per sample) of 16S reads assigned to chloroplasts versus bacteria in individual samples. Reads were grouped into two categories based on taxonomic assignment at the order level (Chloroplast vs Bacteria, with non-chloroplast reads—including missing order assignments—classified as bacteria), and relative abundances were computed within each sample. Samples are shown separately for the two *HaCWINV2* allelic classes (*HaCWINV2*+ and *HaCWINV2*−). Colours indicate read category (chloroplast in green, bacteria in orange), and bar outlines delineate category boundaries within samples.

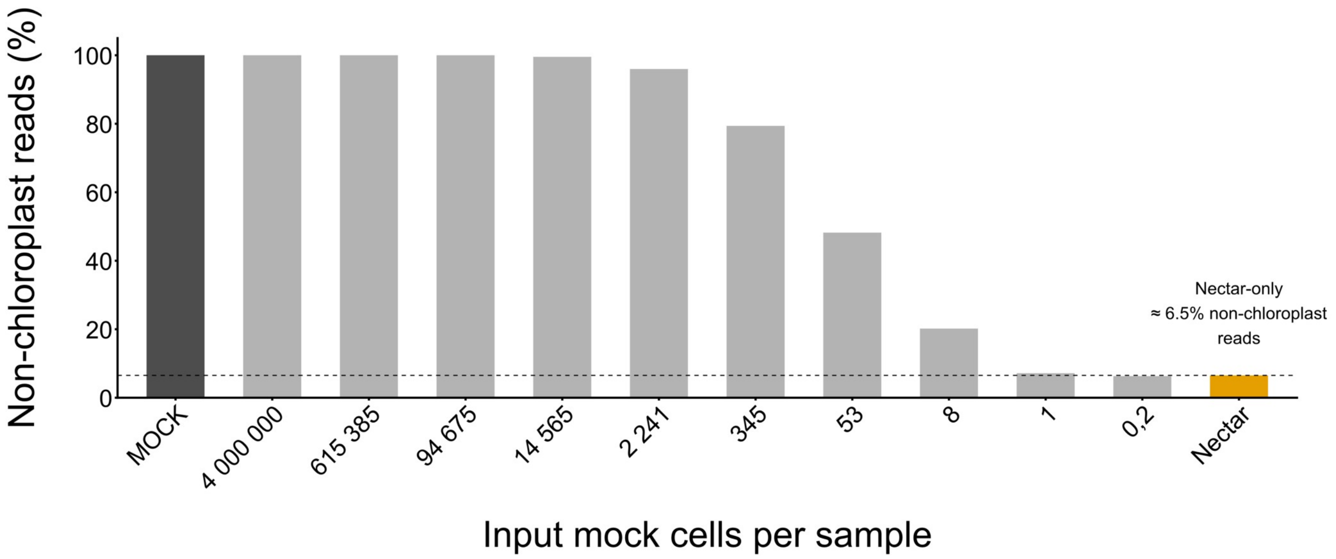


**Supplementary Figure 9. Serial dilution of a bacterial mock community spiked into nectar DNA reveals the detection limit above the nectar background in 16S data.**
A 10-point serial dilution of a bacterial mock community (Zymo; nominal input $2 \times 10^9$ cells/mL) was spiked into a pooled nectar DNA extract, and sequenced alongside a nectar-only control. The x-axis indicates the theoretical number of mock cells per sample (dilution 1: $4.0 \times 10^6$ cells; dilutions 2–10: successive 6.5-fold dilutions), with the undiluted mock and the nectar-only sample shown as reference points. Bars show the proportion of non-chloroplast reads (% of total 16S reads per sample), where non-chloroplastic OTUs were defined as OTUs assigned to bacterial taxa excluding plastids. Bar colours distinguish sample types (undiluted mock, dilution series, nectar-only). The dashed horizontal line indicates the background level of non-chloroplast reads observed in the nectar-only control, used as a practical baseline to infer the limit of detection.

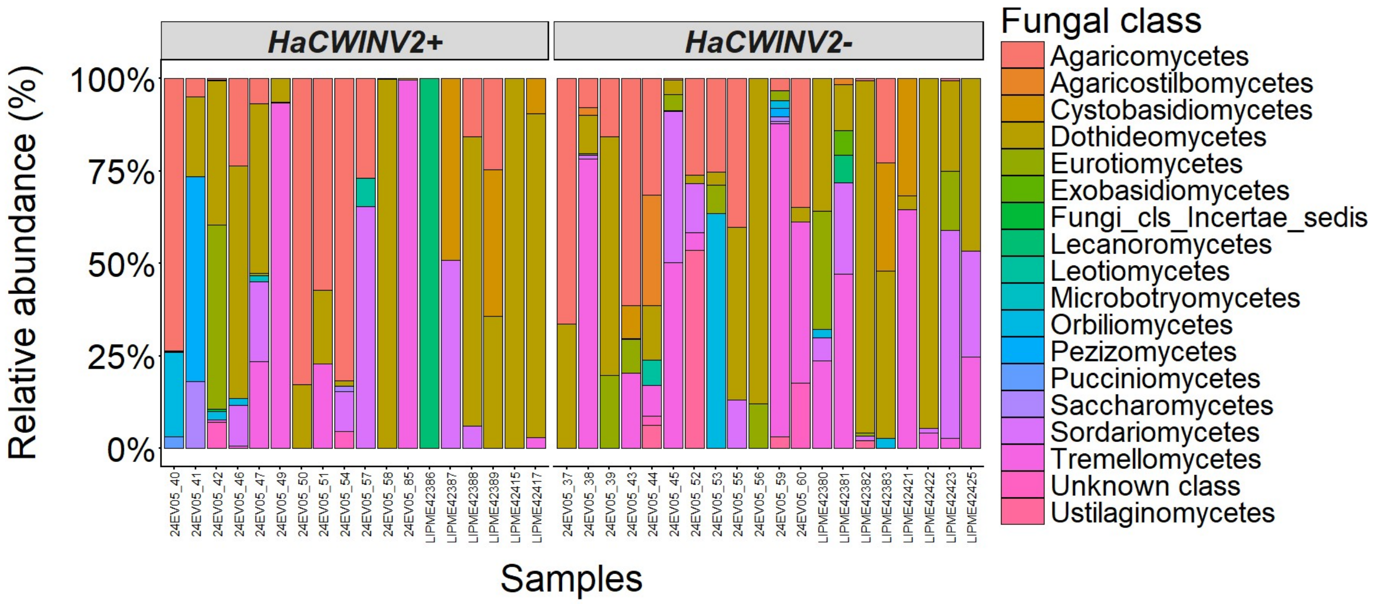


**Supplementary Figure 10. Relative abundance (% of total reads per sample) of fungal taxa in individual nectar samples from sunflower near isogenic lines varying at the *HaCWINV2* locus.** OTUs were grouped at the class level (unclassified taxa shown as Unknown class), and relative abundances were computed within each sample. Samples are shown separately for the two lines (*HaCWINV2*+ and *HaCWINV2*−).

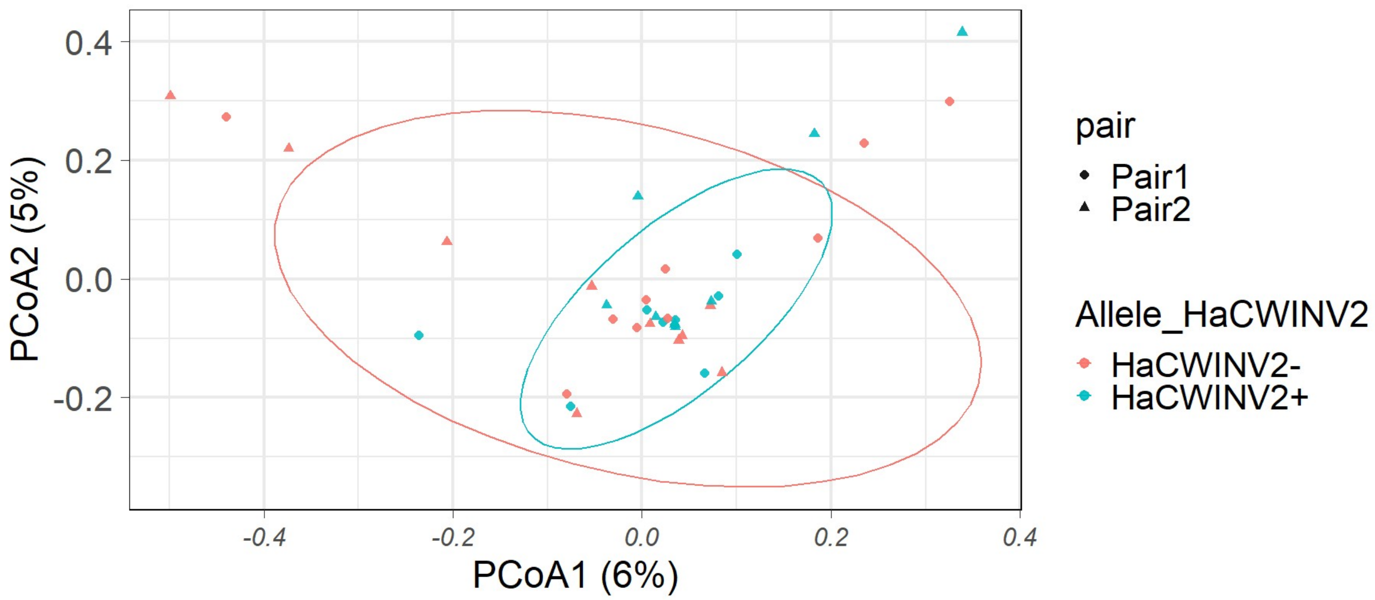


**Supplementary Figure 11. Allele-associated differences in nectar fungal community composition.** Principal coordinates analysis (PCoA) based on Bray–Curtis dissimilarity distances calculated from relative abundances (proportions of total reads per sample) of OTUs. Points represent individual samples, coloured by sunflower line (*HaCWINV2*− and *HaCWINV2*+) and shaped by NIL pair; ellipses indicate 95% confidence intervals around group centroids. The percentage of variance explained by each axis is shown in parentheses. Community dissimilarities were further assessed using PERMANOVA (adonis2) including allele, pair, and run effects, and homogeneity of multivariate dispersions was tested using betadisper (9999 permutations). Sample sizes per NIL pair were: Pair 1, n = 10 plants per allele; Pair 2, n = 8 *HaCWINV2+* plants and n = 10 *HaCWINV2-* plants.

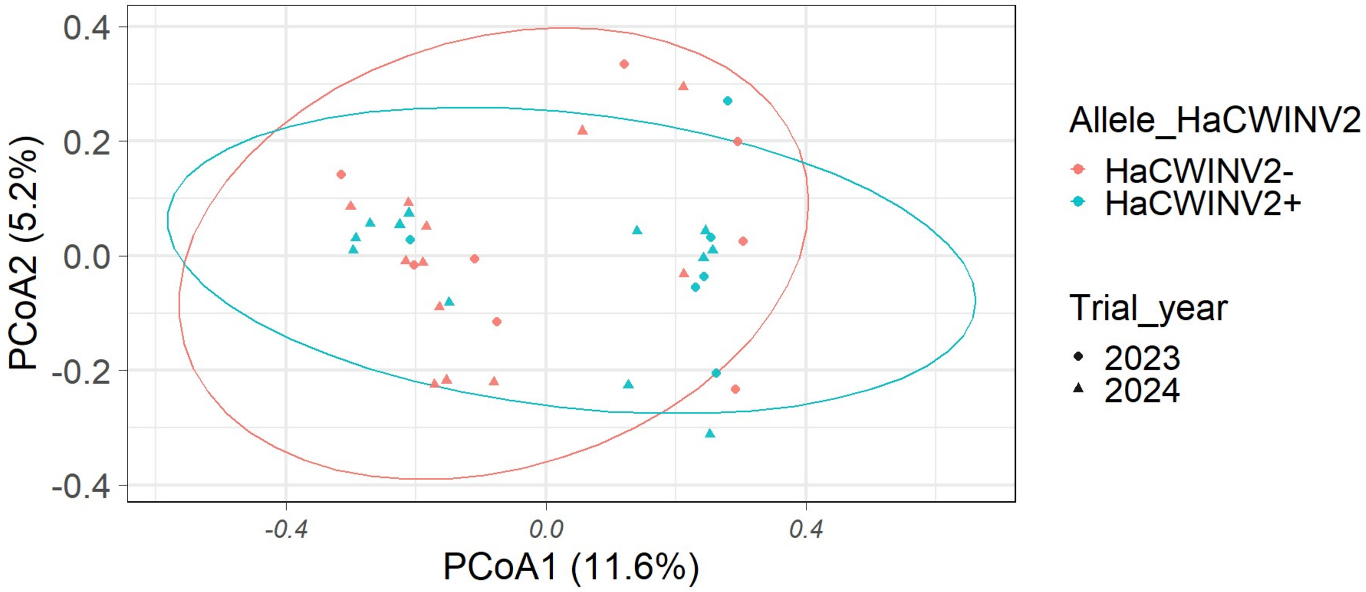


**Supplementary Figure 12. Allele-associated differences in nectar fungal community composition (Jaccard distance).**

Principal coordinates analysis (PCoA) based on Jaccard dissimilarity distances calculated from binary OTU presence –absence data. Points represent individual samples, coloured by sunflower line (*HaCWINV2−* and *HaCWINV2+*) and shaped by NIL pair; ellipses indicate 95% confidence intervals around group centroids. The percentage of variance explained by each axis is shown in parentheses. Community dissimilarities were further assessed using PERMANOVA (adonis2) including allele, pair, and run effects, and homogeneity of multivariate dispersions was tested using betadisper (9999 permutations). Sample sizes per NIL pair were: Pair 1, n = 10 plants per allele; Pair 2, n = 8 *HaCWINV2+* plants and n = 10 *HaCWINV2−* plants.
