## supplementary table for "A plant single nucleotide polymorphism impacts nectar sugar composition, microbial diversity and pollinator visits"

| Gene | SNP position | Ref/ Alt | Allele flanking Primer | Allele-specific Primer 1 | Allele-specific Primer 2 |
|---|---|---|---|---|---|
| HanXRQr2_Chr01g0013961 | HanXRQChr01:64031062..64031062 | C/G | AAGTAGCTAGTTTT-TACGCTTATG | GAAGGTGACCAAGTTCATGCTAATGTGAGATATGAAGAATTAATACGAGG | GAAGGTCGGAGTCAACGGATTT-GTGAGATATGAAGAATTAATACGAGC |
| HanXRQr2_Chr01g0013961 | HanXRQChr01:64031066..64031066 | G/A | TGCAAAATGT-GAGATATGAAGA | GAAGGTGACCAAGTTCATGCTC-TAGTTTTTACGCTTATGCCTCG | GAAGGTCGGAGTCAACGGATTCTAGTTTTTACGCTTATGCCTCA |
| HanXRQr2_Chr01g0013961 | HanXRQChr01:64031362..64031362 | G/T | TGGATGTAGATAC-TACCGAAAACAA | GAAGGTGACCAAGTTCATGCTAACCGAATGGTCAATCAAACTCCT | GAAGGTCGGAGTCAACGGATTCGAATGGTCAATCAAACTCCG |
| HanXRQr2_Chr01g0013961 | HanXRQChr01:64033801..64033801 | A/C | GACATGCA-CATGTGTTCACTTTA | GAAGGTGACCAAGTTCATGCTTCACGACTCGTACCTATATGCTATGTTG | GAAGGTCGGAGTCAACGGATT-GATCACGACTCGTACCTATATGCTATGTTT |
| HanXRQr2_Chr01g0013961 | HanXRQChr01:64034149..64034149 | A/G | GAAAGCCAT-GATTCTTTTGAAG | GAAGGTGACCAAGTTCATGCTGTTTCGTAGAATTGATGTGTTTGTTCC | GAAGGTCGGAGTCAACGGATT-GTTTCGTAGAATTGATGTGTTTGTTCT |
| HanXRQr2_Chr01g0013971 | HanXRQChr01:64037184..64037184 | G/A | GCCAGTTCAAGT-GAACATTTCGTCT | GAAGGTGACCAAGTTCATGC-TAGCCGTTGAAGGGTCACA | GAAGGTCGGAGTCAACGGATT-GCCGTTGAAGGGTCACG |
| HanXRQr2_Chr01g0013971 | HanXRQChr01:64037239..64037239 | G/A | ACTGGCTTCAAGATTGGATTGTTGT | GAAGGTGACCAAGTTCATGCTCCATACCTCCAAGAATGGGTT | GAAGGTCGGAGTCAACGGATTCCATACCTCCAAGAATGGGTC |
| HanXRQr2_Chr01g0013971 | HanXRQChr01:64037435..64037435 | G/A | GCACTCTG-TATCAACGGACATGATT | GAAGGTGACCAAGTTCATGCTGGTACTAGTGCGGGTTTTAACG | GAAGGTCGGAGTCAACGGATT-GGTACTAGTGCGGGTTTTAACA |
| HanXRQr2_Chr01g0013971 | HanXRQChr01:64038889..64038889 | G/C | CA-TAACATGTGGCCACTGCATAATC | GAAGGTGACCAAGTTCATGCTCCATATGATGAGAGCACGACTG | GAAGGTCGGAGTCAACGGATTCCATATGATGAGAGCACGACTC |
| HanXRQr2_Chr01g0013971 | HanXRQChr01:64043601..64043601 | G/A | CAC-CATTGCCATTGAAAGTATTGAA | GAAGGTGACCAAGTTCATGC-TAGGGTTTTGCTATGTTGCATCT | GAAGGTCGGAGTCAACGGATT-GGTTTTGCTATGTTGCATCC |
| HanXRQr2_Chr01g0013971 | HanXRQChr01:64043646..64043646 | G/T | AAGTATTGAAGAT-TATGGGGATGC | GAAGGTGACCAAGTTCATGCTTCATTGGGACAATGGATA | GAAGGTCGGAGTCAACGGATTTCATTGGGACAATGGATC |
| HanXRQr2_Chr01g0013971 | HanXRQChr01:64043854..64043854 | G/A | TGTTTGAA-TATAAACTAAACTTTGCTGGT | GAAGGTGACCAAGTTCATGCTT-GGCAGCATATGGATAAAGTTG | GAAGGTCGGAGTCAACGGATTT-GGCAGCATATGGATAAAGTTA |

**Supplementary Table 1. SNP markers and primers used for PACE genotyping**. For each SNP, the table reports the gene identifier (HanXRQr2 annotation), genomic position on the HanXRQ reference genome (chromosome and coordinate), reference and alternative alleles (Ref/Alt), the common flanking primer, and the two allele-specific primers used for PACE genotyping. Primer sequences are given in the 5′→3′ orientation. Allele-specific primers include standard PACE tails for fluorescent allele discrimination.

| Response | Pair | n (plants) | Shapiro–Wilk normality *P* | Levene (Allele) *P* |
|---|---|---|---|---|
| Nectar volume | Pair1 | 33 | 0.271 | 0.583 |
| Sugar mass | Pair1 | 33 | 0.069 | 0.569 |
| Nectar volume | Pair2 | 40 | 0.274 | 0.419 |
| Sugar mass | Pair2 | 40 | 0.047 | 0.962 |

**Supplementary Table 2. Diagnostic checks for the ANOVA models fitted to plant-level mean nectar volume and sugar mass.** For each response and NIL pair, residual normality was assessed with Shapiro–Wilk tests on model residuals, and homogeneity of variances across *HaCWINV2* alleles was evaluated with Levene's tests. Reported values are p-values.

| Trait | Pair | n | Shapiro–Wilk normality *P* | Levene (Allele) *P* |
|---|---|---|---|---|
| Floret length | Pair1 | 12 | 0,606 | 0,581 |
| Floret width | Pair1 | 12 | 0,284 | 0,438 |
| Floret length | Pair2 | 12 | 0,942 | 0,498 |
| Floret width | Pair2 | 12 | 0,624 | 0,553 |

**Supplementary Table 3. Assumption checks for plant-level mean floret length and width prior to comparisons.** Normality was assessed using Shapiro–Wilk tests and homogeneity of variances using Levene's tests, reported separately for each NIL pair.

| Model | Dispersion (*P*) | Zero inflation ratio (*P*) | Outliers: n / N (*P*) | Uniformity (*P*) | Collinearity: max VIF | Singularity |
|---|---|---|---|---|---|---|
| Bees | 0.426 (0.374) | 1.128 (0.349) | 5 / 9599 (0.14) | 0.023 (5.9e-05) | 4.69 | FALSE |
| Bumblebees | 0.919 (0.841) | 0.992 (0.729) | 2 / 9599 (0.4) | 0.012 (0.137) | 4.52 | FALSE |

**Supplementary Table 4. Diagnostic checks and multicollinearity assessment for the negative-binomial GLMMs (bees and bumblebees).** DHARMa residual diagnostics were computed from 10,000 simulated residuals. Multicollinearity was assessed using variance inflation factors (VIF) from the performance package; singularity was evaluated with check_singularity(). Outlier p-values from testOutliers(..., type = "bootstrap").

| Primers | |
|---|---|
| 16S-tailed_F : | TTTCTGTTGGTGCTGATATTGCAGRGTTYGATYMTGGCTCAG |
| 16S-tailed_R : | ACTTGCCTGTCGCTCTATCTTCRGYTACCTTGTTACGACTT |
| ITS4-tailed_R : | ACTTGCCTGTCGCTCTATCTTCTCCTCCGCTTATTGATATGC |
| ITS1F-tailed_F : | TTTCTGTTGGTGCTGATATTGCTTGGTCATTTAGAGGAAGTAA |

**Supplementary Table 5. Adapter-tailed primer sequences used for long-amplicon nanopore sequencing.** Adapter-tailed primers were used to amplify the bacterial 16S rRNA gene (16S-tailed_F/16S-tailed_R) and the fungal ITS1–ITS4 region (ITS1F-tailed_F/ITS4-tailed_R). Sequences are reported 5′→3′ and include the 5′ adapter tails required for Oxford Nanopore ligation library preparation, followed by the locus-specific primer sequence.

| Metric (response) | Transform used | Shapiro–Wilk normality *P* | Levene (Allele × pair) *P* |
|---|---|---|---|
| Richness | sqrt | 0.238 | 0.892 |
| ENS1 | none | 0.366 | 0.598 |
| ENS2 | none | 0.661 | 0.838 |

**Supplementary Table 6. Diagnostic checks for alpha-diversity analyses.** For each response variable, the transformation used in the final model is indicated. Normality of model residuals was evaluated using Shapiro–Wilk tests. Homogeneity of residual variances was assessed using Levene's tests across HaCWINV2 allele × NIL pair groups. Reported values correspond to p-values.

| Distance | Factor | betadisper permutest *P* |
|---|---|---|
| Bray–Curtis | HaCWINV2 allele | 0.256 |
| Bray–Curtis | pair | 0.209 |
| Bray–Curtis | Trial year | 0.130 |
| Jaccard | HaCWINV2 allele | 0.115 |
| Jaccard | pair | 0.502 |
| Jaccard | Trial year | 0.240 |
| Unweighted UniFrac | HaCWINV2 allele | 0.018 |
| Unweighted UniFrac | pair | 0.812 |
| Unweighted UniFrac | Trial year | 0.381 |

**Supplementary Table 7. Diagnostic assessment of the homogeneity-of-dispersion assumption for beta-diversity analyses**. For each distance metric and grouping factor, multivariate dispersion was tested using betadisper followed by permutest with 9,999 permutations. Reported values are permutation p-values (significant results indicate unequal dispersion among groups).

| Trial | Pair | Allele_*HaCWINV2* | Number of plants | Photos per plant (min –max) | Number of days | Number of pictures in total |
|---|---|---|---|---|---|---|
| 23TO01 | Pair 1 | *HaCWINV+* | 5 | 1212-2868 | 17 | 8718 |
| | | *HaCWINV-* | 5 | 1622-2308 | 18 | 9315 |
| | Pair 2 | *HaCWINV+* | 5 | 1237-2322 | 13 | 8621 |
| | | *HaCWINV-* | 5 | 1025-1886 | 14 | 7165 |
| 24EV05 | Pair 1 | *HaCWINV+* | 4 | 386-1811 | 11 | 5077 |
| | | *HaCWINV-* | 4 | 1343-2305 | 12 | 7489 |
| | Pair 2 | *HaCWINV+* | 5 | 1345-1922 | 13 | 8262 |
| | | *HaCWINV-* | 4 | 677-1730 | 11 | 5371 |
| 23TE21 | Pair 2 | *HaCWINV+* | 6 | 1629-2306 | 13 | 11531 |
| | | *HaCWINV-* | 6 | 1645-2297 | 13 | 11712 |
| 25TE24 | Pair 1 | *HaCWINV+* | 5 | 2005-2839 | 18 | 12201 |
| | | *HaCWINV-* | 5 | 2008-2776 | 18 | 12404 |
| | Pair 2 | *HaCWINV+* | 2 | 1293-2024 | 11 | 3317 |
| | | *HaCWINV-* | 2 | 1693-1883 | 10 | 3576 |

**Supplementary Table 8. Sampling structure of the pollinator dataset across trials, NIL pairs and *HaCWINV2* alleles after daytime and late-flowering filtering.** The table reports the number of individual plants included in the analyses (Number of plants), the range of sampling effort per plant expressed as the minimum and maximum number of photographs (Photos per plant (min–max)), the number of distinct sampling days retained after filtering (Number of days), and the total number of photographs analysed within the corresponding group (Number of pictures in total). Trial indicates the experimental environment (year × field). Pair refers to the near-isogenic line (NIL) pair. Allele_*HaCWINV2* denotes the *HaCWINV2* allelic class (*HaCWINV2*+ or *HaCWINV2*−). Counts are reported after restricting the dataset to daytime photographs only (06:00–21:59) and to the first 99% of each plant's cumulative visitation curve, thereby excluding late-season images with very low visitation and capitula bearing very few florets (typically <10).

| SampleID | 16S_Raw_Reads | 16S_OTU_Raw | 16S_Filtered_Reads | 16S_OTU_Filtered | ITS_Raw_Reads | ITS_OTU_Raw | ITS_Filtered_Reads | ITS_OTU_Filtered |
|---|---|---|---|---|---|---|---|---|
| 24EV05_37 | 49947 | 5 | 0 | 0 | 49569 | 18 | 4166 | 9 |
| 24EV05_38 | 49965 | 2 | 0 | 0 | 49647 | 22 | 13357 | 16 |
| 24EV05_39 | 49962 | 2 | 0 | 0 | 49695 | 12 | 3731 | 7 |
| 24EV05_40 | 49886 | 4 | 0 | 0 | 48765 | 15 | 11709 | 9 |
| 24EV05_41 | 49799 | 5 | 0 | 0 | 49612 | 15 | 35861 | 9 |
| 24EV05_42 | 182 | 1 | 0 | 0 | 48350 | 34 | 47471 | 28 |
| 24EV05_43 | 49971 | 2 | 0 | 0 | 49218 | 27 | 14414 | 17 |
| 24EV05_44 | 49970 | 5 | 0 | 0 | 48934 | 36 | 2158 | 23 |
| 24EV05_45 | 47398 | 93 | 0 | 0 | 48745 | 43 | 19817 | 32 |
| 24EV05_46 | 49921 | 8 | 0 | 0 | 49119 | 32 | 21216 | 15 |
| 24EV05_47 | 49826 | 17 | 0 | 0 | 49109 | 26 | 6106 | 15 |
| 24EV05_49 | 47623 | 10 | 5322 | 5 | 49452 | 19 | 31525 | 11 |
| 24EV05_50 | 49756 | 5 | 0 | 0 | 49421 | 12 | 30096 | 7 |
| 24EV05_51 | 49799 | 6 | 0 | 0 | 49764 | 9 | 9242 | 5 |
| 24EV05_52 | 49792 | 18 | 0 | 0 | 49525 | 26 | 5311 | 16 |
| 24EV05_53 | 49067 | 26 | 22511 | 12 | 49807 | 16 | 36710 | 8 |
| 24EV05_54 | 49642 | 7 | 0 | 0 | 49350 | 15 | 14717 | 8 |
| 24EV05_55 | 49559 | 33 | 1593 | 23 | 49346 | 15 | 7174 | 10 |
| 24EV05_56 | 49766 | 11 | 0 | 0 | 49663 | 15 | 10421 | 8 |
| 24EV05_57 | 49689 | 11 | 0 | 0 | 49239 | 12 | 5031 | 4 |
| 24EV05_58 | 0 | 0 | 0 | 0 | 49595 | 14 | 44260 | 8 |
| 24EV05_59 | 49757 | 27 | 0 | 0 | 49534 | 29 | 32879 | 16 |
| 24EV05_60 | 49916 | 4 | 0 | 0 | 49691 | 18 | 3362 | 9 |
| 24EV05_85 | 49742 | 6 | 0 | 0 | 49551 | 12 | 7446 | 7 |
| LIPME42378 | 5084 | 4 | 0 | 0 | 167 | 1 | 0 | 0 |
| LIPME42379 | 14737 | 8 | 0 | 0 | 20271 | 8 | 0 | 0 |
| LIPME42380 | 4653 | 69 | 1072 | 35 | 12864 | 27 | 9649 | 22 |
| LIPME42381 | 18482 | 14 | 0 | 0 | 12266 | 10 | 9438 | 7 |
| LIPME42382 | 11903 | 11 | 0 | 0 | 43605 | 19 | 32136 | 12 |
| LIPME42383 | 32824 | 14 | 0 | 0 | 15495 | 17 | 1664 | 10 |
| LIPME42384 | 1106 | 9 | 0 | 0 | 14226 | 3 | 0 | 0 |
| LIPME42385 | 22999 | 16 | 0 | 0 | 13056 | 8 | 0 | 0 |
| LIPME42386 | 10809 | 13 | 0 | 0 | 4393 | 3 | 4097 | 1 |
| LIPME42387 | 4860 | 7 | 0 | 0 | 13441 | 7 | 2710 | 3 |
| LIPME42388 | 1299 | 3 | 0 | 0 | 27573 | 10 | 4639 | 5 |
| LIPME42389 | 810 | 2 | 0 | 0 | 14214 | 7 | 1801 | 3 |
| LIPME42414 | 4446 | 5 | 0 | 0 | 1301 | 4 | 0 | 0 |
| LIPME42415 | 16356 | 8 | 0 | 0 | 15621 | 2 | 3864 | 1 |
| LIPME42416 | 7555 | 5 | 0 | 0 | 12473 | 6 | 0 | 0 |
| LIPME42417 | 1671 | 27 | 0 | 0 | 25068 | 13 | 8391 | 8 |
| LIPME42418 | 8501 | 11 | 0 | 0 | 0 | 0 | 0 | 0 |
| LIPME42419 | 958 | 3 | 0 | 0 | 999 | 1 | 0 | 0 |
| LIPME42420 | 14823 | 8 | 0 | 0 | 10750 | 4 | 0 | 0 |
| LIPME42421 | 5431 | 6 | 0 | 0 | 15319 | 15 | 1783 | 9 |
| LIPME42422 | 284 | 1 | 0 | 0 | 4286 | 10 | 2957 | 4 |
| LIPME42423 | 4906 | 24 | 0 | 0 | 8730 | 26 | 5157 | 14 |
| LIPME42424 | 1873 | 4 | 0 | 0 | 1548 | 6 | 0 | 0 |
| LIPME42425 | 11986 | 31 | 0 | 0 | 13678 | 11 | 3317 | 6 |

**Supplementary Table 9. Per-sample sequencing depth and OTU richness before and after contaminant filtering (16S and ITS).** For each sample, the table reports the number of reads and OTUs prior to filtering ("Raw") and after filtering ("Filtered") for both marker datasets. OTU means the number of OTUS per sample and reads means the number of reads per sample. Filtering was performed in R (vegan, ape, phyloseq) and relied on blank and empty-capillary controls: OTUs reaching >5% relative abundance in any negative-control library (after within-control normalization) were removed from the full dataset. OTUs of non-microbial or host origin were discarded (non-fungi ITS sequences in the ITS dataset; chloroplast and mitochondrial sequences in the 16S dataset), and Malasseziales OTUs (human-associated) were removed. Iterative abundance and depth filtering was then applied until convergence by removing OTUs with <10 total reads across samples and samples with <1,000 reads.

| Genotype | Allele_HaCWINV2 | Pair | Plant_ID | Media | Nb_Colonies | Sampling_Date | Counting_Date |
|---|---|---|---|---|---|---|---|
| H03 | HaCWINV2- | Pair2 | 24EV05_31 | R2A | 0 | 05/07/2024 | 09/07/2024 |
| H03 | HaCWINV2- | Pair2 | 24EV05_32 | R2A | 0 | 05/07/2024 | 09/07/2024 |
| H03 | HaCWINV2- | Pair2 | 24EV05_33 | R2A | 1 | 05/07/2024 | 09/07/2024 |
| A03 | HaCWINV2+ | Pair2 | 24EV05_34 | R2A | 0 | 05/07/2024 | 09/07/2024 |
| A03 | HaCWINV2+ | Pair2 | 24EV05_35 | R2A | 0 | 05/07/2024 | 09/07/2024 |
| A03 | HaCWINV2+ | Pair2 | 24EV05_36 | R2A | 1 | 05/07/2024 | 09/07/2024 |
| B06 | HaCWINV2+ | Pair1 | 24EV05_25 | R2A | 6 | 12/07/2024 | 15/07/2024 |
| B06 | HaCWINV2+ | Pair1 | 24EV05_26 | R2A | 2 | 12/07/2024 | 15/07/2024 |
| B06 | HaCWINV2+ | Pair1 | 24EV05_27 | R2A | 5 | 12/07/2024 | 15/07/2024 |
| A06 | HaCWINV2- | Pair1 | 24EV05_28 | R2A | 1 | 12/07/2024 | 15/07/2024 |
| A06 | HaCWINV2- | Pair1 | 24EV05_29 | R2A | 1 | 12/07/2024 | 15/07/2024 |
| A06 | HaCWINV2- | Pair1 | 24EV05_30 | R2A | 2 | 12/07/2024 | 15/07/2024 |
| H03 | HaCWINV2- | Pair2 | 24EV05_31 | TSA | 4 | 05/07/2024 | 09/07/2024 |
| H03 | HaCWINV2- | Pair2 | 24EV05_32 | TSA | 0 | 05/07/2024 | 09/07/2024 |
| H03 | HaCWINV2- | Pair2 | 24EV05_33 | TSA | 1 | 05/07/2024 | 09/07/2024 |
| A03 | HaCWINV2+ | Pair2 | 24EV05_34 | TSA | 2 | 05/07/2024 | 09/07/2024 |
| A03 | HaCWINV2+ | Pair2 | 24EV05_35 | TSA | 3 | 05/07/2024 | 09/07/2024 |
| A03 | HaCWINV2+ | Pair2 | 24EV05_36 | TSA | 0 | 05/07/2024 | 09/07/2024 |
| B06 | HaCWINV2+ | Pair1 | 24EV05_25 | TSA | 25 | 12/07/2024 | 15/07/2024 |
| B06 | HaCWINV2+ | Pair1 | 24EV05_26 | TSA | 9 | 12/07/2024 | 15/07/2024 |
| B06 | HaCWINV2+ | Pair1 | 24EV05_27 | TSA | 11 | 12/07/2024 | 15/07/2024 |
| A06 | HaCWINV2- | Pair1 | 24EV05_28 | TSA | 1 | 12/07/2024 | 15/07/2024 |
| A06 | HaCWINV2- | Pair1 | 24EV05_29 | TSA | 8 | 12/07/2024 | 15/07/2024 |
| A06 | HaCWINV2- | Pair1 | 24EV05_30 | TSA | 3 | 12/07/2024 | 15/07/2024 |

**Supplementary Table 10. Culturability of nectar-associated microbes on semi-selective media (2024).** Nectar microbial culturability was assessed in 2024 by plating nectar samples onto tryptic soy agar (TSA) and R2A agar. For each plant, nectar was diluted in 500 µL sterile phosphate-buffered saline (PBS), spread-plated using sterile beads, and incubated for minimum 3 days at 28 °C, after which colony-forming units (CFUs) were counted. The table reports, for each plant (Plant_ID) within each NIL pair and HaCWINV2 allelic class (HaCWINV2+ or HaCWINV2−), the growth medium used (TSA or R2A), the number of colonies counted (Nb_Colonies; CFUs per plate), and the sampling and counting dates.
