## supplementary methods for "A plant single nucleotide polymorphism impacts nectar sugar composition, microbial diversity and pollinator visits"

### Variant identification and filtering on the cultivated *H. annuus* "SEAM" population

The Sunflower Extended Association Mapping population, hereafter called "SEAM", is an extended version of the Sunflower Association Mapping population, which is referred to in the literature as "SAM" population and which was originally presented in Mandel *et al*., 2013. The 281 accessions of sunflower inbred lines found in the "SAM" include restorer and maintainer lines, branched and unbranched lines, as well as oil and non-oil lines. Among them, 29 originate from the INRAE Genetic Resource Centre for Sunflower and Soybean (INRAE Toulouse) and make part of the core collection presented in Coque *et al.* (2008) and exploited in Cadic *et al*. (2013).

An additional number of 154 accessions were added within the frame of the AGRI4POL project. Sequencing libraries for these accessions were prepared using the Illumina DNA Prep kit (Illumina, San Diego, CA, USA) and the MGIEasy Universal Library Conversion kit (MGI, Shenzhen, Guangdong, China). Subsequently, the libraries were sequenced in 2x150 bp paired-end mode using the DNBSeq-T7RS High troughput Sequencing Kit (MGI) and the DNBSEQ-T7 sequencer (MGI), and aiming for a target depth of 10X per accession (Get-Plage sequencing facility, INRAE Toulouse). Eventually, the "SEAM" population included therefore 435 accessions.

To identify genomic variations and build the polymorphism matrix, raw sequencing reads were first pre-processed for quality using Trimmomatic v 0.39 (Bolger et al 2014) to remove adapter sequences and low-quality bases. The following parameters were imposed: trimmo_params = '-validatePairs -phred33', trimmo_leading_qual_min = 20, trimmo_trailing_qual_min = 20, trimmo_sliding_window_size = 4, trimmo_sliding_window_qual = 20, trimmo_min_length = 50. Trimmed reads were aligned to the sunflower reference genome HanXRQr2.0 (GCA_002127325.2) using BWA-MEM v 0.7.17 (Li 2013). To ensure high-quality alignments, the resulting BAM files were filtered by retaining only properly paired reads and removing PCR duplicates with SAMtools v 1.9 (Danecek et al 2021). Additionally, genomic regions with a coverage depth > 100X were excluded to minimize biases arising from repetitive sequences using SAMtools v 1.9 (Danecek et al 2021). Final sequencing depth ranged from 6.1X to 57.4X, with an average value of 10.9X.

Initial variant discovery (SNVs and INDELs) was performed for each genotype individually using the 'mpileup2snp' and 'mpileup2indels' functions of VarScan v 2.4.4 (Koboldt et al 2012). The analysis was run with the following parameters: varscan_mutant_min_coverage = 5, varscan_mutant_min_reads2 = 3, varscan_mutant_avg_qual = 20, varscan_mutant_var_freq = 0.2, varscan_mutant_var_freq_for_hom = 0.8, varscan_mutant_pvalue = 0.01. To generate a comprehensive polymorphism matrix, all identified variant positions were consolidated into a unified set of loci. Genotypes were then re-interrogated at these specific positions using the 'mpileup2cns' function of VarScan v 2.4.4 (Koboldt et al 2012) to ensure consistent calling across the entire population, including the recovery of reference alleles. The individual results were then merged into a final global matrix. Finally, the coding effect of the identified polymorphisms was predicted using SnpEff v 5.1 (Cingolani et al 2012). The .vcf file obtained after these steps included a total number of 88,863,075 SNVs and 11,569,647 INDELs.

Variant filtering was then realized as follows. First, INDELs and multiallelic SNPs were filtered out. Subsequently, 16 individuals presenting a percentage of total missing genotype calls > 20% were removed. At this stage, the dataset included therefore 419 individuals. Afterwards, four additional steps of filtering were applied, namely by removing: (i) SNPs with DP < 10 and DP > 50; (ii) SNPs with GQ < 25; (iii) SNPs presenting missing values in > 20% of the genotypes; (iv) SNPs mapping to scaffolds and cpDNA. All the aforementioned steps were performed using VCFtools (Danecek et al 2011).

After these filtering steps, missing values were imputed using Beagle v 5.5 (Browning et al 2018) with standard parameters. Subsequently, SNPs presenting MAF < 0.05 were discarded. Eventually, the thus obtained .vcf file contained a total number of 1,842,523 SNPs.

### Variant identification on the wild *H. annuus* "Heliawild" population

The “Heliawild” population included a total number of 184 wild *H. annuus* accessions, namely 148 accessions newly sequenced within the frame of study and 36 accessions previously sequenced within the frame of the SUNRISE and described in Badouin *et al.*, 2017. All these accessions are maintained at the INRAE Genetic Resource Centre for Sunflower and Soybean (INRAE Toulouse), and have been selected based on geographical collection data available through the USDA National Plant Germplasm System (https://www.grin-global.org).

Sequencing libraries for the newly sequenced accessions were prepared using the TruSeq DNA Nano kit (Illumina) and sequenced in 2x150 bp paired-end mode using an AVITI sequencer (Element Biosciences, San Diego, CA, USA), and aiming for a target depth of 20X per accession (Get-Plage sequencing facility, INRAE Toulouse).

The identification of variants on the “Heliawild” population was performed as described in the case of the “SEAM” population and, after the steps of trimming and alignment, the sequencing depth ranged from 9.2X to 27.5X, with an average value of 13.5X.

The filtering steps, instead, were not carried out in this case. A .vcf file containing the initially discovered (*i.e.* not filtered) 341,513,778 variant positions (SNVs and INDELs) for the 184 *H. annuus* accessions is available at https://www.heliagene.org/HeliaWild/index.html.

**Insect detection model : training dataset, augmentation pipeline, and performance evaluation**

Images were compiled from experiments conducted over a three-year period and consisted of time-series acquisitions collected during flowering to monitor insect visitation. The model was trained to detect three classes: bees, bumble bees, and moths/butterflies. Insects were annotated as polygons, excluding transparent wing regions as well as legs and antennae, to reduce annotation inconsistency when image quality was insufficient to delineate these structures. Image quality was heterogeneous, ranging from high-quality images to degraded images affected by blur, out-of-focus acquisition, environmental conditions such as morning dew, or miscalibrated active lighting during night-time imaging.

To prevent data leakage arising from temporal background continuity within image series, the dataset was split into training, validation, and test sets while enforcing that all images from a given time series belonged to the same partition. Empty images were added in the train partition to reduce overfitting to recurrent backgrounds.

Native image resolutions were 1188 × 2112, 1692 × 3008, 2160 × 3840, 2376 × 4224, and 3420 × 6080 pixels. To preserve object resolution while limiting computational cost, each training data sample was divided into multiple 1024 × 1024 crops randomly centered on each insect, while ensuring that no insect appeared more than once across crops. Validation and test images were kept at native dimensions to enable a reliable estimation of false positive detections in backgrounds.

The final training set comprised 6,124 images (768 × 768 and 1024 × 1024), including 1,196 empty images, with 2,982 bee, 675 bumble bee, and 1,613 moth/butterfly annotations.
The validation set comprised1,715 images at mixed native resolutions, with no empty images, with 1,066 bee, 201 bumble bee, and 650 moth/butterfly annotations.
The test set comprised 1340 images at mixed native resolutions, with no empty images, with 886 bee, 148 bumble bee, and 426 moth/butterfly annotations.

Modelling was performed using the YOLO11x architecture through our public library deepvisiontools (https://forge.inrae.fr/ue-apc/librairies/python/deepvisiontools) wrapper built on Ultralytics (https://github.com/ultralytics/ultralytics). Instance masks were converted to bounding boxes after augmentation.
Standard data augmentations were applied using torchvision, including color transformations, rotations and flips, blur, and scale jitter. To further reduce dependence on redundant backgrounds, a custom augmentation was added in which image backgrounds were replaced, with a probability of 15%, by backgrounds sampled from the public Kaggle Flowers dataset

(https://www.kaggle.com/datasets/imsparsh/flowers-dataset), while preserving insect masks and their associated pixels.

After optimization of the confidence and non-maximum suppression thresholds, the performance of this model and dataset version (04-2025) is reported in the public GitLab repository.

**Supplementary References**


- Bolger AM, Lohse M, Usadel B (2014). Trimmomatic: a flexible trimmer for Illumina sequence data. Bioinformatics, Aug 1;30(15):2114-20
- Cadic E, Coque M, Vear F, Grezes-Besset B, Pauquet J, Piquemal J, Lippi Y, Blanchard P, Romestant M, Pouilly N, Rengel D, Gouzy J, Langlade N, Mangin B, Vincourt P (2013). Combined linkage and association mapping of flowering time in Sunflower (Helianthus annuus L.). Theor Appl Genet 126, 1337–1356
- Coque M, Mesnildrey S, Romestant M, Grezes-Besset B, Vear F, Langlade NB, Vincourt P (2008). Sunflower nested core collections for association studies and phenomics, in: Proceeding of the 17th Int. Sunflower Conference. Córdoba, Spain: International Sunflower Association. pp. 725–28 https://hal.science/hal-05527727v1.
- Li H (2013). Aligning sequence reads, clone sequences and assembly contigs with BWA-MEM. arXiv preprint arXiv:1303.3997
- Danecek P, Bonfield JK, Liddle J, John Marshall, Ohan V, Pollard MO, Whitwham A, Keane T, McCarthy SA, Davies RM, Li H (2021). Twelve years of SAMtools and BCFtools. GigaScience, vol 10(2)
- Koboldt DC, Zhang Q, Larson DE, Shen D, McLellan MD, Lin L, Miller CA, Mardis ER, Ding L, Wilson RK (2012). VarScan 2: somatic mutation and copy number alteration discovery in cancer by exome sequencing. Genome Research, Mar;22(3):568-76
- Cingolani P, Platts A, Wang LL, Coon M, Nguyen T, Wang L, Land SJ, Lu X, Ruden DM (2012). A program for annotating and predicting the effects of single nucleotide polymorphisms, SnpEff: SNPs in the genome of Drosophila melanogaster strain w1118; iso-2; iso-3. Fly (Austin), Apr-Jun;6(2):80-92
- Danecek P, Auton A, Abecasis G, Albers CA, Banks E, DePristo MA, Handsaker R, Lunter G, Marth G, Sherry ST, McVean G, Durbin R and 1000 Genomes Project Analysis Group (2011). The Variant Call Format and VCFtools, Bioinformatics
- Browning BL, Y Zhou Y, and Browning SR (2018). A one-penny imputed genome from next generation reference panels. Am J Hum Genet 103(3):338-348
